# Transient epithelial mimicry reconciles stemness and regional specification in neural crest cells of avian beaks

**DOI:** 10.1101/2025.05.29.656909

**Authors:** Carmen Sánchez Moreno, Alexander V. Badyaev

## Abstract

Multicellular morphogenesis must balance organismal cohesion with local tissue differentiation. Migratory stem cells commonly fulfill these dual needs by orchestrating region-specific tissue differentiation, yet how they balance the maintenance of stemness with positional sensitivity is unclear. Here, we show that in the developing avian beak, early arriving neural crest-derived mesenchymal (NCM) cells transiently match protein profiles of the overlying epithelium, which diverges as development proceeds. This “local mimicry” phase propagates region-specific epithelial signaling into the mesenchyme, producing distinct boundaries that anchor mesenchymal cell condensations. As NCM cells accumulate, cells within condensations undergo morphological and molecular homogenization, erasing regional differences in protein expression and restoring cellular multipotency. These cycles of transient specialization and homogenization – driven by universal processes of cell proliferation and migration – enable NCM cells to reconcile location-specific anchoring signals with stemness needed for ongoing regional specification of growing beak. By balancing global coordination with local divergence, this developmental organization can facilitate the remarkable evolutionary diversification of avian beaks.

## Introduction

In multicellular development, cell differentiation can only occur in the context of cell condensations, where close contact among cells synchronizes and channels their variability and primes fate transitions (Hall 1978; Atchley and Hall 1991; Tacchetti et al. 1992; Dunlop and Hall 1995; Hall 2003; Soldatov et al. 2019). Yet, how initially uniform cells aggregate in the first place and what guides the spatial and temporal distribution of the condensations remain poorly understood. Crucially, it is unclear how these two aspects are reconciled: condensations can be an emergent outcome of aggregation and adhesion of individual cells (Cottrill et al. 1987; Widelitz et al. 1993; Barna and Niswander 2007; Glimm et al. 2014; Kaul et al. 2015; Svandova et al. 2020), yet they have defined spatial and temporal distribution implying positional sensitivity (Wu et al. 2006; Abramyan and Richman 2015; Hu et al. 2015b). This combination of universality and specificity is particularly striking for condensations composed of distantly induced multipotent cells that travel through diverse signaling environments before arriving at diverging local contexts where they form region-specific aggregations (Hall 1980; Gitton et al. 2010; Buitrago-Delgado et al. 2015; Kaucka et al. 2016; Hall 2018; Woronowicz and Schneider 2019; Hovland et al. 2022; Paudel et al. 2022). Mechanisms that modulate the spatial and temporal sensitivity of these cells, and thus reconcile their stemness with their ability to organize locally appropriate tissues, are central to the evolution of morphological diversity.

A particularly well studied example is craniofacial morphogenesis in vertebrates, where migrating neural crest-derived mesenchymal (NCM) cells that build the upper and lower jaws combine remarkable regulatory autonomy with an equally striking ability to orchestrate region-specific molecular and cellular interactions (Hall 1999; Santagati and Rijli 2003; Schneider and Helms 2003; Creuzet et al. 2005; Jheon and Schneider 2009; Welsh and O’Brien 2009; Minoux and Rijli 2010; Hu et al. 2015b; Fonseca et al. 2017; Selleri and Rijli 2023; Schneider 2024). This combination has fueled a long-standing research program addressing whether spatio-temporal specificities are embedded in NCM cells at the time and place of their delamination, accumulate in NCM cell groups during migration, or emerge through cellular and molecular interactions with local tissues (Hall 1999; Minoux and Rijli 2010; Kaucka et al. 2016; Schneider 2018b; Selleri and Rijli 2023). Classical grafting experiments, particularly in birds, found evidence for all of these processes although with notable differences across NCM cell sites or origin (Noden 1983; Richman and Tickle 1989; Trainor et al. 2002; Hu et al. 2003; Creuzet et al. 2005; Hu and Marcucio 2012; Hall et al. 2014; Ray and Chapman 2015). Essentially, migrating NCM cells, although retaining substantial molecular memory of their origin and accumulating migration bias *en route* (some of which is coopted for establishing growth polarity) still retain enough multipotency upon their arrival to orchestrate region-specific condensations (Buitrago-Delgado et al. 2015; Dupin et al. 2018; Kelsh et al. 2021). Importantly, the epithelial-mesenchymal signaling that is essential for proper placement and orientation of condensations is remarkably non-specific to either regions within beaks or to beaks themselves but instead is deeply evolutionarily conserved (Eames and Helms 2004; Le Douarin et al. 2004; Schneider 2005; Lu et al. 2024), which raises the question of how location-specificity emerges from the interaction of two non-location-specific players – migrating stem cells and universal signaling for their anchoring. A particular challenge in this process is the acquisition of sequential positional identity without compromising multipotentiality of NCM cells – local specializations should not bias subsequent developmental repertoire.

Mechanisms capitalizing on predictable material properties of cells and tissues (such as tension or elasticity), as well as on measures of distance, time, and compartmentalization are particularly well suited for such non-biasing positional regulation, including in NCM tissues (Francis-West et al. 1998; Medeiros and Crump 2012; Barriga et al. 2018; Meinecke et al. 2018; Ransom et al. 2018; Lenne and al. 2021; Badyaev et al. 2025a). One potential resolution therefore is if the distribution of condensations is linked to the distance and timing of their cell’s migration and to associated changes in either sensitivity to epithelial signaling or epithelial signaling itself caused by developmental expansion. Indeed, the ability of epithelial signaling to initiate mesenchymal condensation is time-sensitive, NCM cell fate specification is affected by the accumulated molecular bias of migration or origin, and expanding epithelium invariably results in a feed-forward signaling gradient (Hall and Miyake 1992; MacDonald and Hall 2001; Haworth et al. 2004; Liu et al. 2005; Merrill et al. 2008; Welsh and O’Brien 2009; Cela et al. 2016). Thus, the important step in establishing how universal signaling succeeds in delivering stem cells to proper locations is to establish the time, place, and contexts in which these cells first acquire location-specificity and to identify how time- and distance-dependent local divergence can be integrated with conserved epithelial-mesenchymal signaling.

Here we take this approach by tracing the onset of positional identity in cellular and molecular determinants of NCM cell condensations across the beak primordia of the house finch (*Haemorhous mexicanus*). We specifically focus on formation of the condensation boundary. The boundary delineates condensations within the fields of homogeneous mesenchymal cells and is essential for condensation growth, modulation of external inputs, maintenance of uniformity and ultimate differentiation (Hall and Miyake 2000; Giffin et al. 2019). Whether the condensation boundary is a cause or consequence of condensation formation is unclear; it can emerge as a passive consequence of cell aggregation and resulting radial signaling gradient, or it can precede condensations through upregulation of adhesion signaling in mesenchymal cells in interaction with extracellular matrix components, producing physical or signaling barriers of compacted cells facilitating sorting and retention of migrating mesenchymal cells (Widelitz et al. 1993; Hall and Miyake 2000; Giffin et al. 2019). Here, we explore an additional explanation that combines these processes (Simsek and Özbudak 2022; Kicheva and Briscoe 2023): that early-arriving NCM cells amplify or propagate local epithelial signaling into the mesenchyme thus forming signaling or cellular boundaries that seed regional condensations.

We evaluate these scenarios by first establishing the temporal sequence of tissues reorganization involved in condensation formation (Figs. 1, S2) and the contribution of cellular and molecular mechanisms to this reorganization (Fig. 2). We then capitalize on extensive developmental divergence and tissue reorganization within and among the upper and lower jaw primordia (Depew and Compagnucci 2008; Gitton et al. 2010; Tak et al. 2017) to infer the importance of the global and local molecular and cellular processes for condensation formation and subsequent reprograming and local specification (Figs. 1, 2).

**Figure 1.**
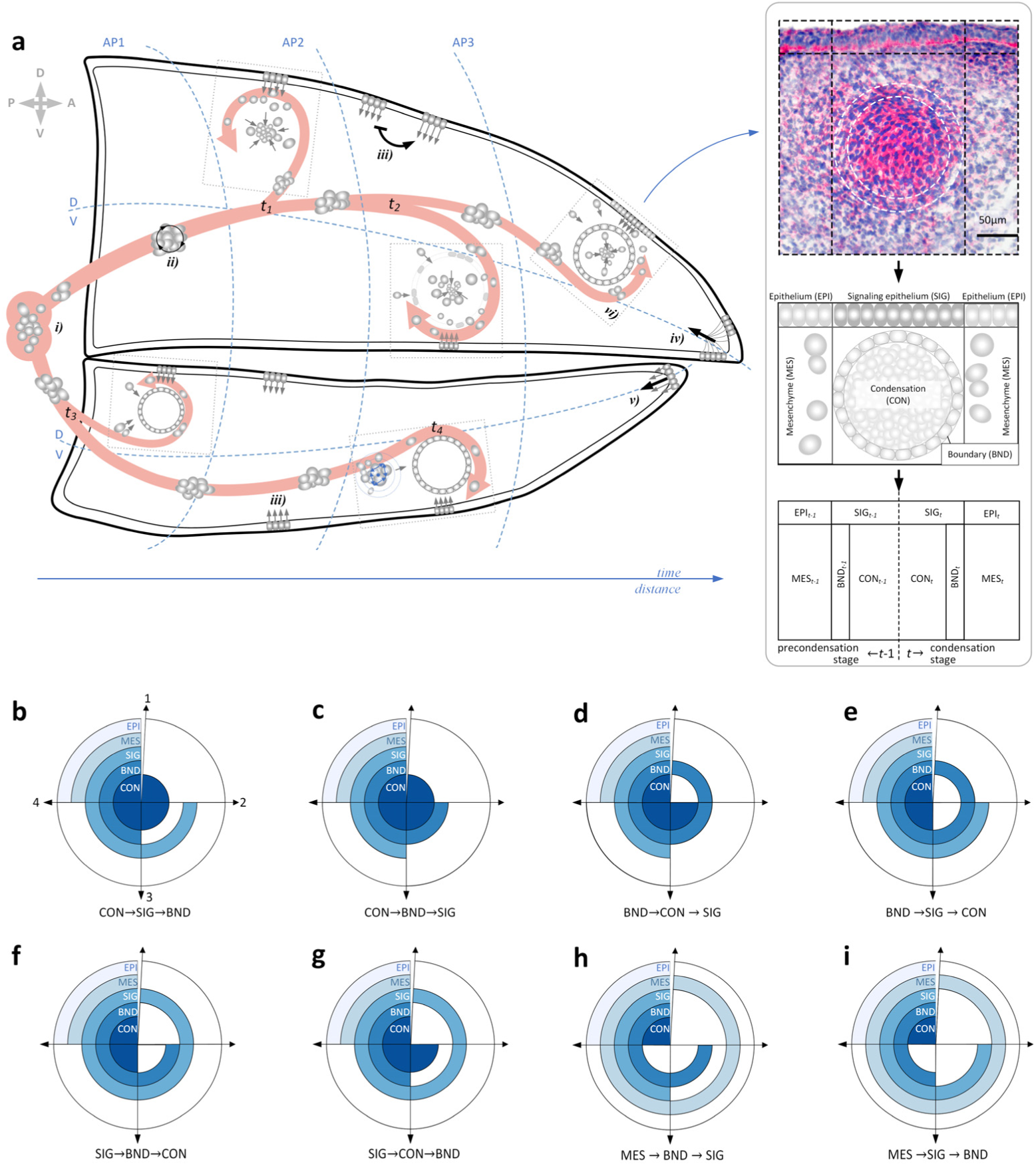
Reconciling stemness and position-specificity in the origin of condensations. **a)** Schematics of NCM cell migration streams (coral arrows) moving in groups from their induction sites in neural folds (*i*) to the future condensations distributed across anterior-posterior (AP) and dorsoventral (DV) zones of the upper and lower beaks. En route, groups of migrating NCM cells interact with each other (*ii*), experience external input, such as unfolding feed-forward epithelial signaling (*iii*) and axial signaling gradients emanating from the base and tip of beaks (e.g., “boundary signaling” (frontonasal ectodermal zone, *iv*) and consolidated signaling (e.g., “hinge-cap” model, *v*)). Dashed rectangles at condensation sites represent grid boxes analyzed in this study (see Methods). Insert (*vi*) shows an example of Ihh expression (magenta) in studied tissues (see also Fig. S2) and corresponding tissue classification within each grid box: CON – condensation, BND – boundary, MES – uncondensed mesenchyme, SIG – signaling epithelium overlying condensation, and EPI – adjacent epithelium. Also shown is the corresponding tissue scheme for reporting results for pre- (*t-1*) and condensation (*t*) stages. **(b-i)** known and proposed temporal sequences of condensation origin. **(b-c)** arriving NCM cells either induce local epithelium signaling (b) or proliferate, forming condensation and its boundary, subsequently inducing epithelial signaling (c); **(d-e)** compaction of earlier arriving NCM cells forms a boundary that sorts or traps cells either producing condensations directly (d) or inducing epithelial signaling that facilitates condensation formation (e), **(f-g)** signaling epithelium either induces the formation of a boundary leading to accumulation of arriving NCM cells (f) or attracts migrating cells causing them to aggregate and form the condensation directly (g). **(h-i)** dynamically interacting NCM cells acquire rigidity and either form a boundary (h) or express proteins enabling them to induce or respond to local epithelial signaling (i).

**Figure 2.**
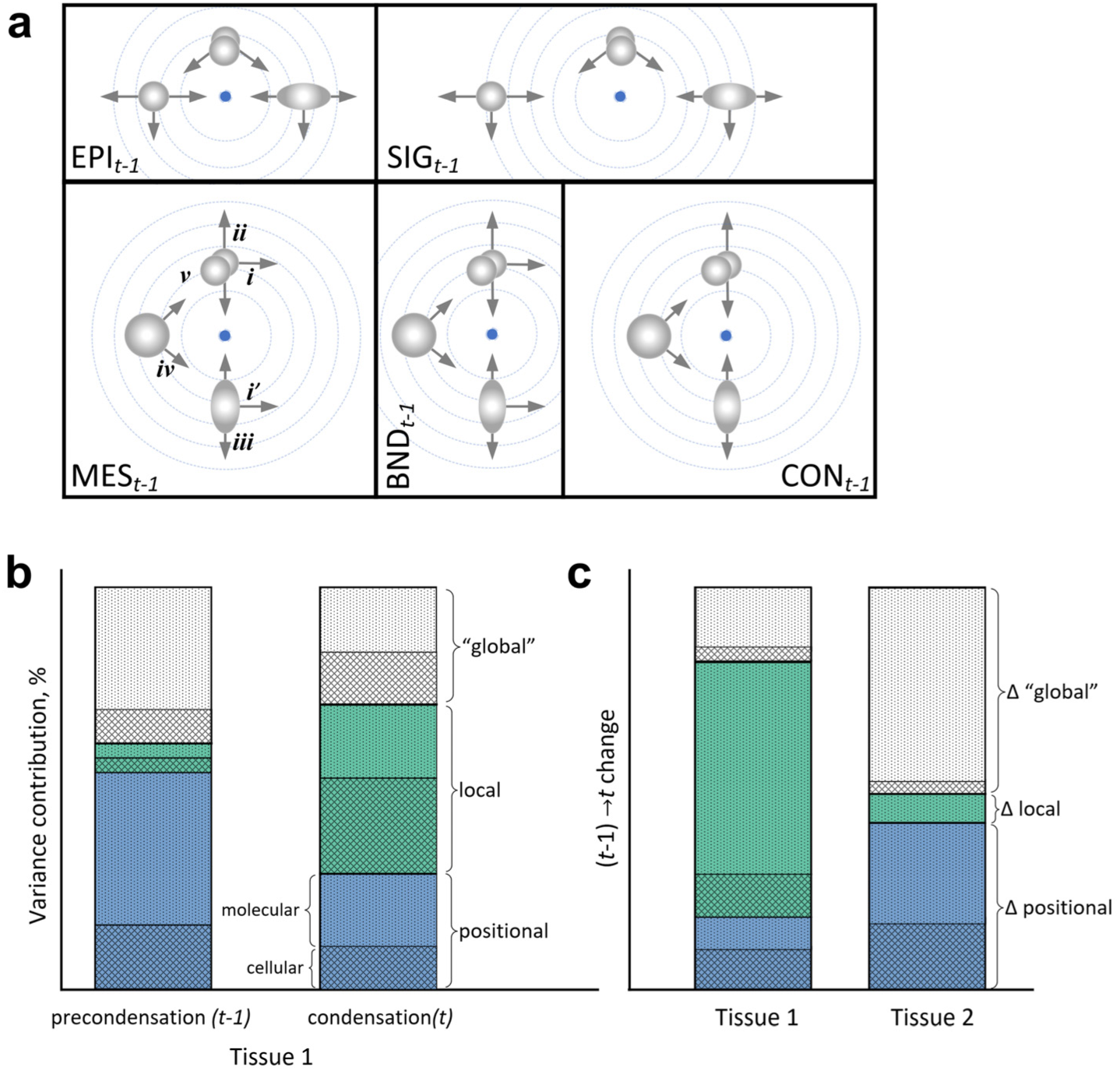
Cellular and molecular processes of condensation formation and inference into their location-specificity. **(a)** Cells can show polarity towards the sites of future condensations (*i*; *i’-* shows variation in cell polarity), towards (*ii*) or away (*iii*) from epithelial or mesenchymal cells, forming cell groups (*iv*). These movements along with changes in cell density and proliferation (*v*) can transport, dilute, or delimit spread of molecular signaling (dashed circles) leading to compartmentalization of condensation sites. Tissue abbreviations as in Fig. 1. Comparison of cell morphology and protein expression in each tissue at different stages across scenarios of Fig. 1 (b-i) informs how multipotent NCM cells orchestrate position-specific condensations. **(b** and **c)** Variance partitioning in molecular and cellular mechanisms to infer tissue reprograming and specification during condensation formation. Predictions illustrating an approach to distinguish “local”, context-dependent variance (contribution of interaction terms between condensation placements along AP and DV axes and between jaws), “positional”, gradient-based variance (contribution of condensation placement along AP and DV axes or in different jaws), and “global” variance unexplained by any location- or jaw-based measure, either directly or in interaction, and potentially representing tissue-specific variation or unmeasured effects (see Methods). Hypothetical contributions of molecular and cellular mechanisms are shown. (b) Variance partitioning between pre- (*t*-1) and condensation (*t*) stages. (c) Change between stages for two tissues. Tissue 1 shows an increase in local specificity, especially of molecular contributors, such as those associated with greater developmental specification, and lessening contribution of positional gradients in cellular and molecular effects. Tissue 2 shows diminishing location-specificity, expected under tissue reprogramming in which positional effects no longer explain significant variance. Increased contribution of positional effects reflects tissue reorganization, here also implying interdependence of condensations due to their temporal synchrony or juxtaposition.

Using an analytical approach that allows us to trace millions of cells throughout avian beak development across the full span of the upper and lower beak primordia (Table 1, Fig. S1), we find that condensations are initiated by the formation of signaling boundaries – NCM cells extending location-specific protein profiles of regional overlying epithelium into the mesenchyme – followed by the accumulation and compaction of NCM cells within these boundaries and subsequent morphological and molecular homogenization. Arriving pre-condensation NCM cells accumulate significant positional- and location-specific variation in their protein profiles and cellular arrangements, but this variation is progressively erased as condensations grow, even as condensations and their boundaries become morphologically distinct. These cycles of transient specialization and homogenization in migratory NCM cells can enable position-specific implementation of a conserved epithelial-mesenchymal signaling without compromising NCM cell ability to seed region-specific condensations. Such developmental organization also reconciles global coordination with local specialization within structures, allowing for the recombination of regulatory modules, thus underpinning evolutionary diversification of avian beaks.

**Table 1.** Sample sizes for condensation-containing grid boxes (*n*_boxes_ =35,755; *n*_cells_= 2.12 x 10^6^) across tissue types during (*t*) and prior (*t-1*) to condensation formation across tissue types, beak prominences, future tissue types (F_Tissue) and upper and lower beak (Jaw).

| Jaw | Prominence | F_Tissue | Condens | Condensation tissues ($t$) | | | | | Tissues prior to condensation formation ($t-1$) | | | | |
| --- | --- | --- | --- | --- | --- | --- | --- | --- | --- | --- | --- | --- | --- |
|  |  |  |  | BND<sub>t</sub> | CON<sub>t</sub> | EPI<sub>t</sub> | MES<sub>t</sub> | SIG<sub>t</sub> | BND<sub>t-1</sub> | CON<sub>t-1</sub> | EPI<sub>t-1</sub> | MES<sub>t-1</sub> | SIG<sub>t-1</sub> |
| Upper | Frontonasal | cartilage | U2 | 204 | 250 | 540 | 435 | 125 | 12 | 13 | 25 | 54 | 14 |
|  |  | cartilage | U5 | 274 | 220 | 371 | 606 | 24 | 20 | 33 | 20 | 91 | 17 |
|  |  | muscle | U8 | 276 | 85 | 462 | 397 | 28 | 68 | 85 | 90 | 238 | 17 |
|  | Lateral nasal | cartilage | U4 | 384 | 452 | 333 | 608 | 38 | 31 | 84 | 20 | 153 | 1 |
|  |  | cartilage | U10 | 81 | 108 | 99 | 288 | 1 | 25 | 50 | 27 | 134 | 23 |
|  |  | cartilage | U12 | 234 | 185 | 260 | 429 | 12 | 26 | 73 | 38 | 131 | 17 |
|  | Maxillary | cartilage | U6,7 | 274 | 334 | 688 | 1,060 | 17 | 93 | 330 | 165 | 495 | 81 |
|  |  | muscle | U9 | 112 | 38 | 677 | 447 | 40 | 36 | 53 | 78 | 155 | 39 |
| Lower | Mandibular | cartilage | L1 | 531 | 221 | 625 | 661 | 34 | 7 | 30 | 21 | 42 | 10 |
|  |  | cartilage | L3,6 | 518 | 187 | 1,257 | 488 | 77 | 191 | 161 | 289 | 308 | 90 |
|  |  | cartilage | L5 | 294 | 33 | 679 | 590 | 0 | 55 | 94 | 83 | 306 | 15 |
|  |  | cartilage | L7 | 504 | 201 | 645 | 449 | 21 | 44 | 39 | 72 | 125 | 14 |
|  |  | muscle | L8 | 824 | 253 | 824 | 1,156 | 26 | 107 | 102 | 122 | 250 | 35 |
|  |  | muscle | L9 | 301 | 26 | 657 | 654 | 18 | 96 | 126 | 60 | 252 | 18 |
|  |  | bone | L2 | 532 | 219 | 627 | 670 | 12 | 9 | 38 | 22 | 43 | 7 |
|  | Meckel's | cartilage | L4 | 467 | 121 | 1,344 | 818 | 8 | 27 | 56 | 57 | 104 | 5 |

## Results

### Matched signaling of mesenchymal boundary and overlying epithelium precedes condensation formation

Although not yet morphologically distinct from surrounding mesenchyme, future boundary cells had elevated Ihh expression compared to all other tissues, and also had distinct expression of β-catenin, Bmp4, Dkk3, and Tgfβ2 relative to adjacent mesenchyme, as well as different Fgf8 and β-catenin expression compared to the overlying epithelium (Fig. 3a, Tables 2, S4). Protein profiles of future boundary cells were more similar to the overlying epithelium than to the adjacent mesenchyme (Fig. 4a). Future condensation cells expressed lower β-catenin compared to both the overlying epithelium and the future boundaries, but did not differ in expression of other proteins or cell morphology from the surrounding mesenchyme (Fig. 3, Table S4). Prior to condensation formation, the epithelium overlying future condensations was similar to the adjacent epithelium in protein expression and cell morphology (Fig. 4). Outside of future condensation sites, mesenchymal tissues showed polarity towards the sites of future condensations whereas future boundary cells showed strong polarity towards the overlying epithelium (Fig. 5a). Mesenchymal and epithelial tissues had distinct protein profiles (except for Ihh and CaM) and differed in cell morphology and variability (Figs. 3, 5, Tables 2, S4).

**Figure 3.**
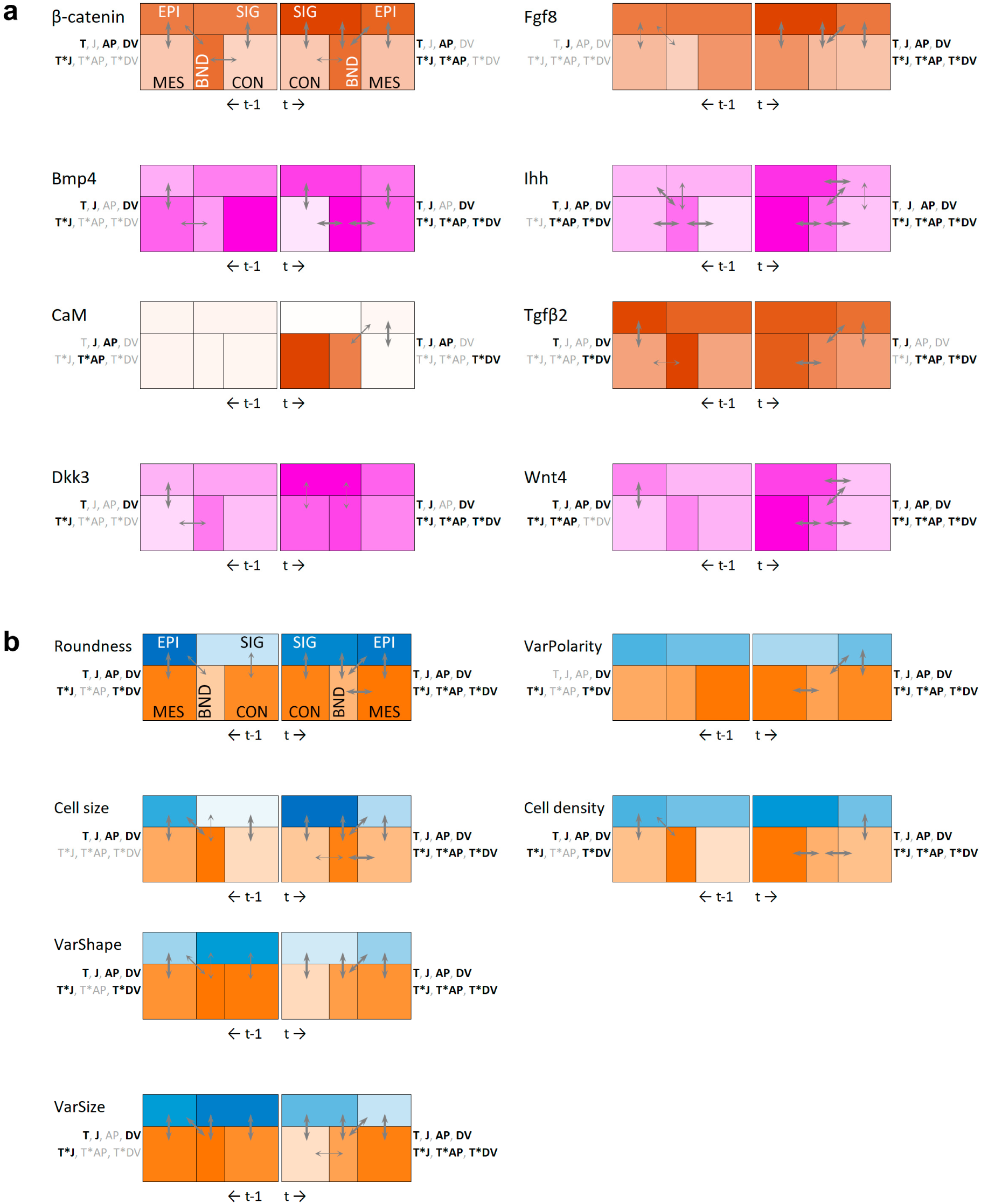
Tissue-specific **(a)** protein expression and **(b)** cell morphology during pre- (*t*-1, left side) and condensation (*t,* right side) stages. Schematics of tissues follows insert of Fig. 1a. Shown are LS means and significant differences between them (based on Tables 2 and S4) as well as associated dependency on tissue type (T), jaw (J), anterior-posterior (AV) or dorsoventral (DV) zones, as well as on interaction between the tissue type and jaw (T*J), anterior-posterior (T*AP) or dorsoventral (DV) axes (based on Table 2). Bold values indicate significance after adjustment for multiple pairwise comparisons. Arrows indicating the LS mean differences are proportional to adjusted *P*-value (thin arrows: *P* ≤ 0.1, medium: 0.05≤ *P* < 0.1, and thick arrows: *P* < 0.01, based on Table S3). Color saturation is proportional to values for all tissues and time periods; color scheme reflects IHC staining group in (a) and is separate for mesenchymal (MES, BND, CON) and epithelium (EPI, SIG) tissues in (b).

**Figure 4.**
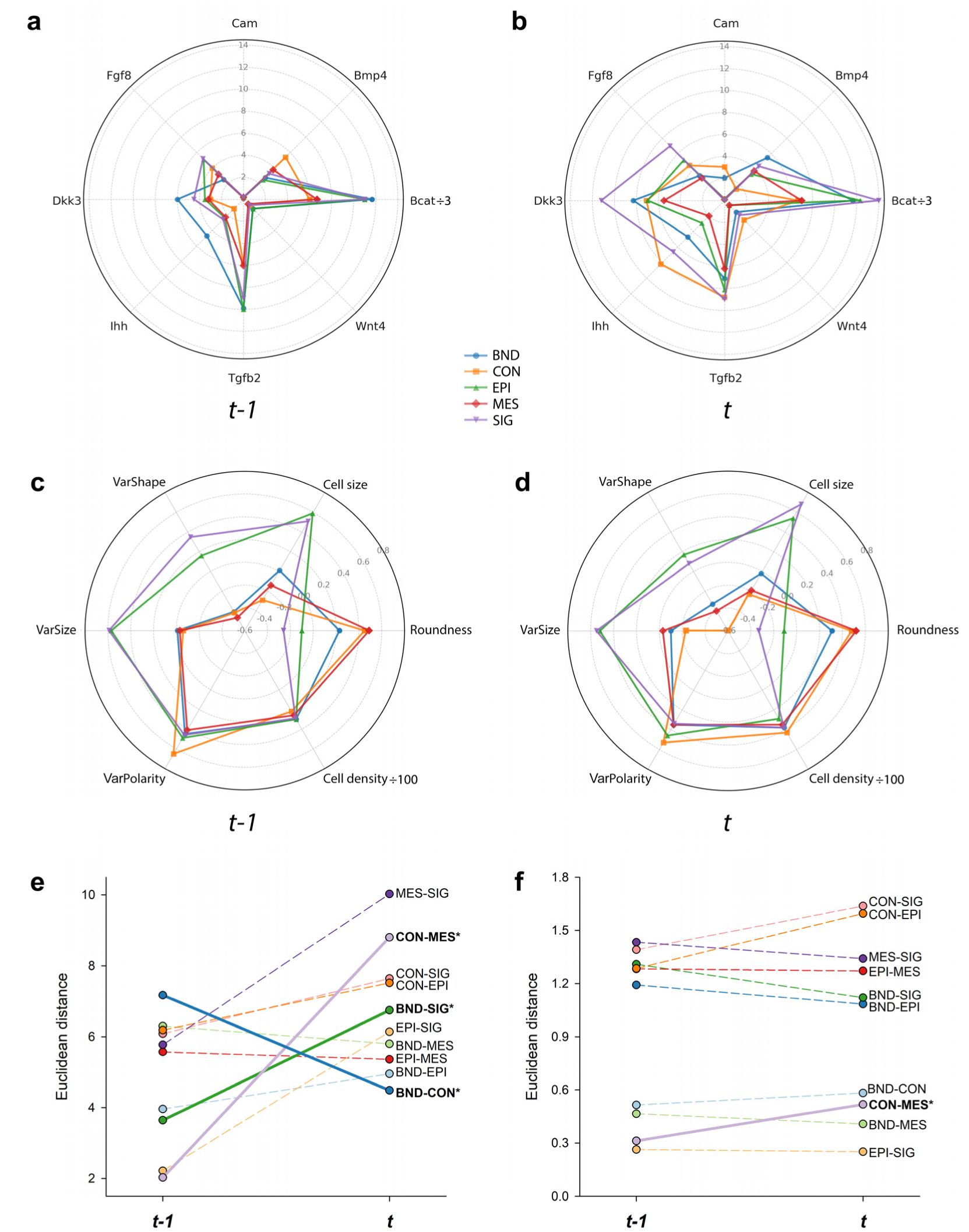
Summary of tissue differences in **(a-b)** protein expression (β-catenin/3 is shown) and **(c-d)** cell morphology (cell density/100 is shown) as well as changes in pairwise tissue distances in **(e)** protein expression and **(f)** cell morphology between pre- (*t-1*) and condensation (*t*) stages (based on Fig. 3). Bold values, solid lines, and asterisks indicate significant change in Euclidean distances (two-tailed *t*-test).

**Figure 5.**
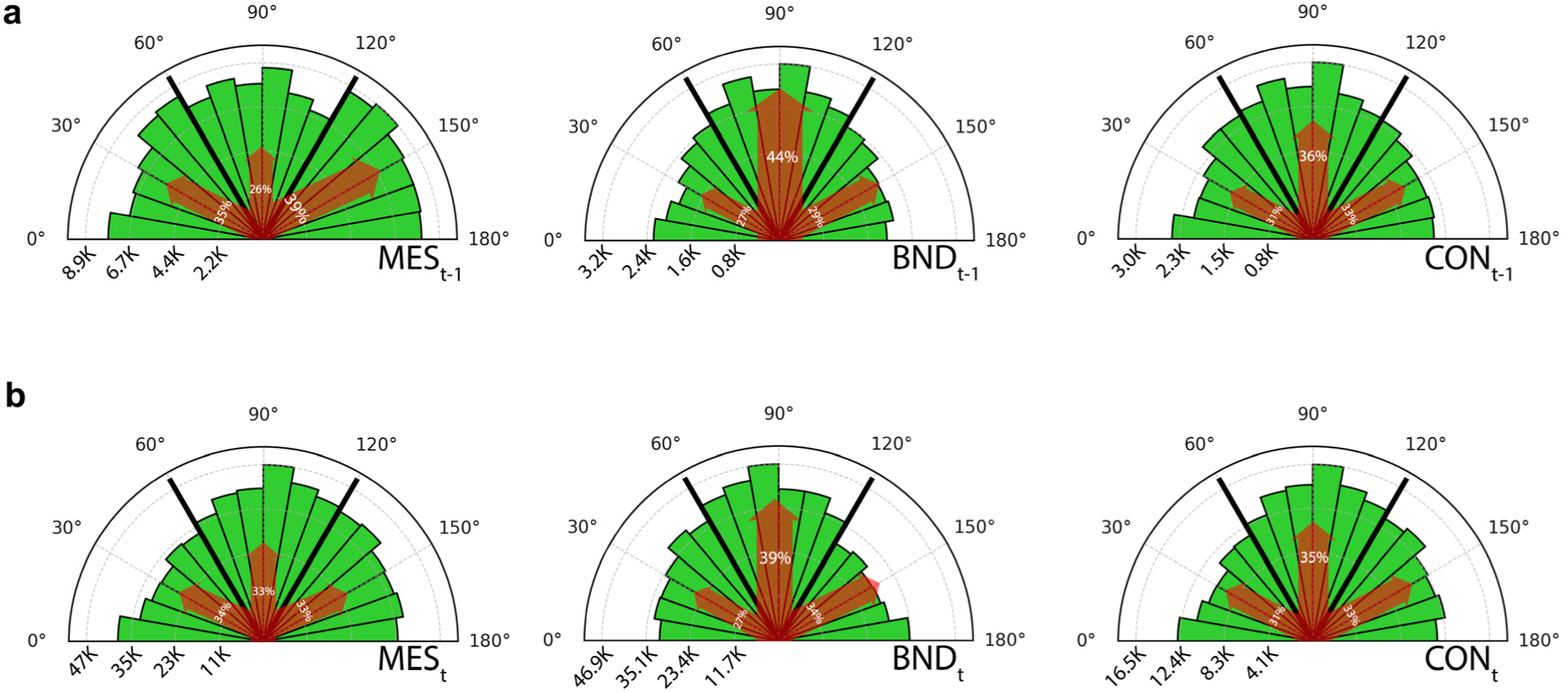
Cell polarity in mesenchymal tissues **(a)** prior (*t*-1) and **(b)** during (*t*) condensation stage. 60-120° tertile encompasses alignments toward or away from the adjacent epithelium. 0-60° and 120-180° tertiles encompasses polarities towards or away from adjacent mesenchymal tissues (1/2 of the combined frequency should be used for direct comparison with the 60-120° tertile frequency). The width of red arrows is proportional to the percentage of cells in the tertile. In precondensation stage (*t-1*), only boundary (BND) and mesenchyme (MES) differ from each other (*F*_2,18K_= 4.1, *P* = 0.02, BND vs MES, *P* = 0.014, Bonferroni-corrected ɑ=0.016). During condensation stage (*t*), the polarity of all tissues differs from each other (*F*_2,18K_=8.6, *P* < 0.001, all pairwise *P*’s <0.01).

### Enclosed condensations become highly distinct from adjacent tissues

Enclosed condensations contained smaller, more densely packed, and more uniform cells compared to the surrounding boundary, and expressed lower β-catenin and Bmp4, but higher Ihh, Wnt4, and Tgfβ2 (Figs. 3ab, 4bd, 5b, Table S4). Condensed and uncondensed mesenchyme differed in the expression of Bmp4, Fgf8, Ihh, Tgfβ2, and Wnt4; condensations contained cells that were more uniform in size and shape and were more densely packed than adjacent uncondensed mesenchyme (Figs. 3, 4d, 5; Table S4). Cells of condensation boundaries were bigger, more oblong in the direction of overlying epithelium (Fig. 3b, 4d, 5b), and distinct from cells of adjacent mesenchyme, particularly in size and polarity (Fig. 4f) and in protein expression (Figs. 3b, 4be; Table S4). The epithelium overlying condensations expressed higher Ihh and Wnt4 than the adjacent epithelium; uncondensed mesenchyme and its overlying epithelium differed in expression of all proteins, except for Dkk3 and Wnt4 (Figs. 3, 4b, Tables 2, S4).

### Growing condensations undergo cellular and molecular homogenization

Changes in molecular and cellular variation during condensation formation were mostly confined to condensed mesenchyme and its overlying epithelium (Fig. 6, Table 3, S5). Bmp4 that was highly expressed at the site of future condensations became non-detectable there, whereas expression of all other proteins increased strongly within condensations and the overlying epithelium (except for Tgfβ2 and CaM that only increased within the condensation) once a condensation was formed (Fig. 6, Table 3, S5). β-catenin and Bmp4 also increased in the epithelium overlying uncondensed mesenchyme while Tgfβ2 and Wnt4 decreased; Dkk3 lso became highly expressed in uncondensed mesenchyme (Fig. 6, Table S5). Within condensations, cells became smaller and more packed, more uniform in size and shape, and less polarized (Figs. 3, 5, 6). Protein profiles of boundaries and condensations converged to each other while diverging from adjacent mesenchyme and epithelium (Figs. 4a,b,e). Similarly, condensations became morphologically distinct from adjacent mesenchyme (Fig. 4f).

**Figure 6.**
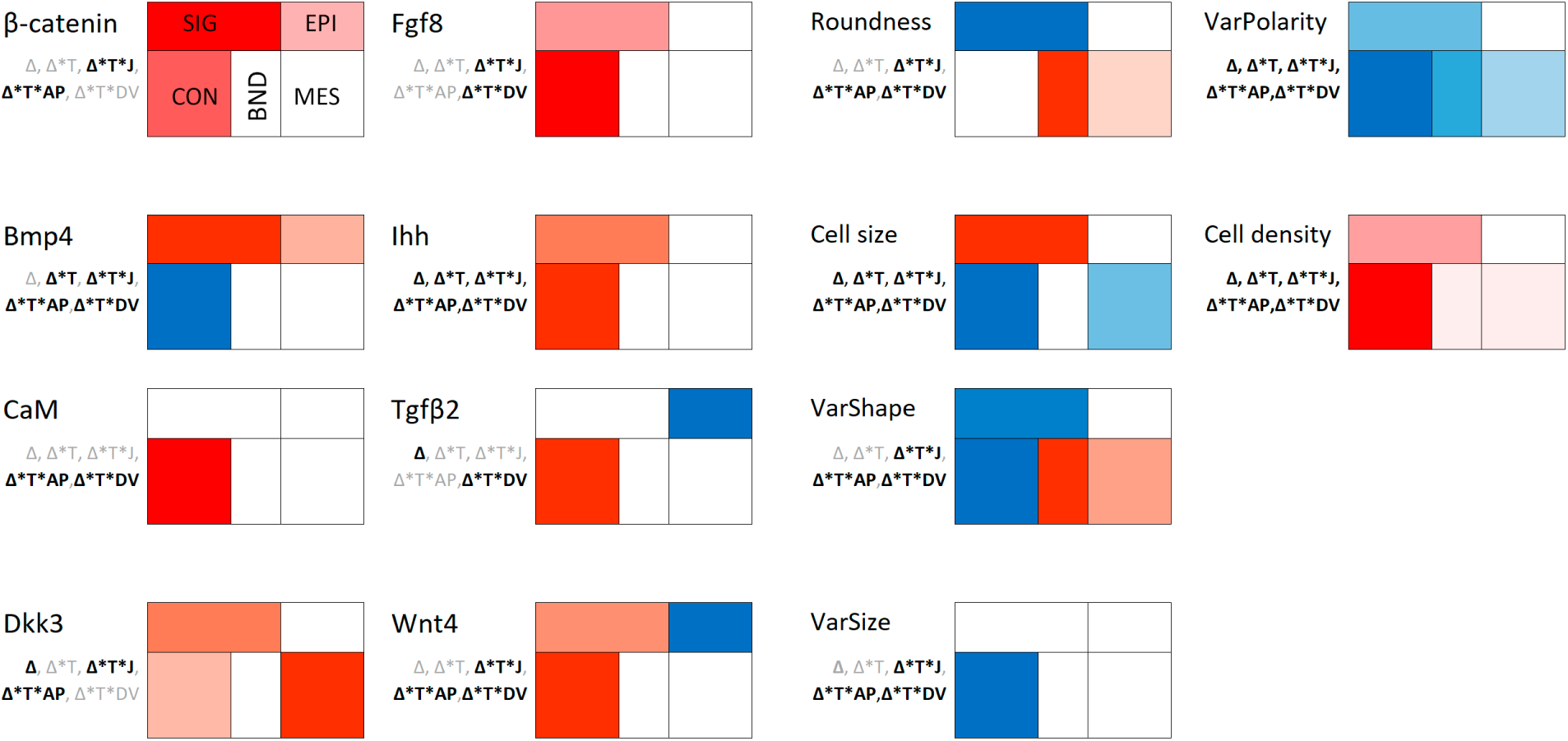
Relative change [(after-before)/before] in protein expression and cell morphology between pre- (*t* -1) and condensation (*t*) stages (data in Table S5). Shown are LS means controlling for the effects of axial position, jaw, and population. Color saturation shows magnitude of change (blue shows negative values, red – positive values) among tissues within each protein or cell measure. Absence of color indicates a lack of significant change. Text shows significance of change (Δ) and its association with tissue type (Δ*T), jaw (Δ*T*J), anterior-posterior (Δ*T*AV) or dorsoventral (Δ*T*DV) axes. Bold values indicate significance at *α* <0.05 (Table 3).

### Axial gradient and location-specificity in condensation formation

Molecular and cellular changes during condensation formation were distinct between locations within the jaw and along anterior-posterior and dorsoventral axes (Fig. 3, Table 3). Prior to condensation formation, β-catenin, CaM, Ihh, Wnt4 and cell morphology, variability (except in polarity and size), and density formed a pronounced anterior-posterior gradient in both jaws, while β-catenin, Bmp4, Dkk3, Ihh, Tgfβ2, Wnt4, and cell morphology, variability and density formed a dorsoventral gradient (Fig. 3, Table 2). Bmp4, CaM, Fgf8, Ihh, and Wnt4, along with cell morphology, density, and variability (except polarity), also differed strongly between upper and lower beak tissues prior to condensation formation (Table 2). Once a condensation was formed, cell morphology and variability depended on condensation location along the anterior-posterior and dorsoventral axes, whereas many proteins showed positional variation along anterior-posterior (except for Bmp4, Dkk3, and Tgfβ2) and dorsoventral (except for β-catenin, CaM and Tgfβ2) axes and between jaws (except for β-catenin) (Table 2).

**Table 2.** Protein expression and cell morphology in relation to tissue type, upper vs lower jaw, anterior-posterior and dorsoventral zones and interactions among these factors prior (t-1) and during (t) condensation formation. Shown are Type III *F* values for fixed effects with variance components covariance structure and Kenward-Roger degrees of freedom from a mixed-effects model and associated *P*-values. Bold values are significant at *P* < 0.05

| Factor | Time | Effects |  |  |  |  |  |  |  |  |  |  |  |  |  |
| --- | --- | --- | --- | --- | --- | --- | --- | --- | --- | --- | --- | --- | --- | --- | --- |
|  |  | Tissue |  | Jaw |  | ZoneAP |  | ZoneDV |  | Tissue*Jaw |  | Tissue*ZoneAP |  | Tissue*ZoneDV |  |
| | | $F$ | $P$ | $F$ | $P$ | $F$ | $P$ | $F$ | $P$ | $F$ | $P$ | $F$ | $P$ | $F$ | $P$ |
| β-catenin | t | **171.2** | <0.01 | 0.1 | 0.71 | **16.5** | <0.01 | 0.2 | 0.81 | **2.4** | 0.05 | **4.3** | <0.01 | 1.8 | 0.07 |
|  | t-1 | **6.8** | <0.01 | 0.30 | 0.59 | **4.7** | <0.01 | **9.4** | <0.01 | **3.1** | 0.02 | 1.0 | 0.44 | 0.5 | 0.81 |
| Bmp4 | t | **11.7** | <0.01 | **4.6** | 0.03 | 2.1 | 0.08 | **6.5** | <0.01 | **2.5** | 0.04 | **4.2** | <0.01 | **5.7** | <0.01 |
|  | t-1 | **3.8** | <0.01 | **4.3** | 0.04 | 0.6 | 0.65 | **7.4** | <0.01 | **3.0** | 0.02 | 0.9 | 0.53 | 1.9 | 0.06 |
| CaM | t | **4.2** | <0.01 | **15.2** | <0.01 | **3.0** | 0.02 | 0.9 | 0.39 | 0.3 | 0.90 | 1.5 | 0.11 | **3.6** | <0.01 |
|  | t-1 | 1.0 | 0.40 | **17.8** | <0.01 | **7.1** | <0.01 | 1.9 | 0.15 | 0.8 | 0.56 | **2.6** | <0.01 | 0.2 | 0.97 |
| Dkk3 | t | **2.9** | 0.02 | **6.4** | 0.03 | 0.7 | 0.62 | **4.1** | 0.02 | **5.5** | <0.01 | **2.6** | <0.01 | **3.0** | <0.01 |
|  | t-1 | **2.5** | 0.04 | 3.1 | 0.15 | 2.1 | 0.09 | **6.8** | <0.01 | **4.5** | <0.01 | 1.1 | 0.36 | 1.2 | 0.31 |
| Fgf8 | t | **37.0** | <0.01 | **14.1** | <0.01 | **7.5** | <0.01 | **9.3** | <0.01 | **6.9** | <0.01 | **2.1** | 0.01 | **4.5** | <0.01 |
|  | t-1 | 1.9 | 0.10 | **6.3** | 0.02 | 0.6 | 0.64 | 0.0 | 0.99 | 0.3 | 0.87 | 1.4 | 0.18 | 1.5 | 0.14 |
| Ihh | t | **16.20** | <0.01 | **15.3** | <0.01 | **16.8** | <0.01 | **9.1** | <0.01 | **6.7** | <0.01 | **2.9** | <0.01 | **3.5** | <0.01 |
|  | t-1 | **2.9** | 0.02 | **17.1** | <0.01 | **6.1** | <0.01 | **9.7** | <0.01 | 0.2 | 0.94 | **3.2** | <0.01 | **2.2** | 0.02 |
| Tgfbβ2 | t | **26.7** | <0.01 | **8.0** | 0.01 | 0.6 | 0.68 | 1.2 | 0.30 | 1.2 | 0.31 | **1.9** | 0.02 | **2.3** | 0.02 |
|  | t-1 | 1.7 | 0.15 | 1.0 | 0.34 | 0.9 | 0.44 | **3.4** | 0.04 | 1.4 | 0.23 | 1.3 | 0.21 | **2.2** | 0.03 |
| Wnt4 | t | **10.4** | <0.01 | **13.6** | <0.01 | **9.4** | <0.01 | **10.3** | <0.01 | **3.7** | <0.01 | **5.0** | <0.01 | **9.6** | <0.01 |
|  | t-1 | **6.0** | <0.01 | **12.0** | <0.01 | **2.4** | 0.05 | **5.8** | <0.01 | **2.4** | 0.05 | **2.3** | 0.00 | 1.6 | 0.13 |
| Roundness | t | **217.2** | <0.01 | **29.4** | <0.01 | **5.01** | <0.01 | **6.5** | <0.01 | **65.6** | <0.01 | **3.7** | <0.01 | **17.4** | <0.01 |
|  | t-1 | **15.5** | <0.01 | **282.7** | <0.01 | **5.5** | <0.01 | **10.4** | <0.01 | **13.6** | <0.01 | 1.6 | 0.07 | **2.0** | 0.04 |
| Size | t | **347.6** | <0.01 | **390.2** | <0.01 | **59.3** | <0.01 | **37.0** | <0.01 | **34.6** | <0.01 | **8.9** | <0.01 | **20.2** | <0.01 |
|  | t-1 | **31.7** | <0.01 | **75.7** | <0.01 | **2.4** | 0.05 | **53.8** | <0.01 | 1.3 | 0.26 | 1.5 | 0.09 | 1.3 | 0.25 |
| VarShape | t | **226.5** | <0.01 | **796.5** | <0.01 | **10.6** | <0.01 | **6.5** | <0.01 | **592.2** | <0.01 | **1.9** | 0.02 | **27.6** | <0.01 |
|  | t-1 | **12.3** | <0.01 | **259.4** | <0.01 | **4.5** | <0.01 | **9.8** | <0.01 | **20.7** | <0.01 | 1.3 | 0.17 | **4.5** | <0.01 |
| VarSize | t | **323.7** | <0.01 | **666.2** | <0.01 | **11.8** | <0.01 | **11.3** | <0.01 | **213.4** | <0.01 | **8.0** | <0.01 | **28.1** | <0.01 |
|  | t-1 | **27.5** | <0.01 | **117.9** | <0.01 | 2.2 | 0.07 | **31.9** | <0.01 | **9.3** | <0.01 | 1.4 | 0.12 | 1.8 | 0.07 |
| VarPolarity | t | **23.4** | <0.01 | **41.6** | <0.01 | **42.8** | <0.01 | **102.8** | <0.01 | **4.8** | <0.01 | **9.9** | <0.01 | **14.2** | <0.01 |
|  | t-1 | 0.3 | 0.88 | 0.00 | 0.98 | 1.6 | 0.16 | **18.1** | <0.01 | **5.1** | <0.01 | **1.5** | 0.09 | **2.0** | 0.04 |
| Density | t | **57.1** | <0.01 | **340.1** | <0.01 | **58.0** | <0.01 | **68.5** | <0.01 | **123.8** | <0.01 | **7.2** | <0.01 | **23.2** | <0.01 |
|  | t-1 | **11.3** | <0.01 | **162.2** | <0.01 | **23.1** | <0.01 | **10.1** | <0.01 | **5.5** | <0.01 | 1.1 | 0.40 | **4.8** | <0.01 |

### Accumulation and erasure of location-specificity in protein profiles during condensation formation

At the precondensation stage, proteins in the overlying epithelium and in the mesenchyme of future condensation boundaries had the most location-specific expression; Dkk3, CaM, Ihh, Fgf8, and β-catenin had the most positional dependency whereas Dkk3, β-catenin, and Wnt4 also had the most location-specific expression (Figs. 7a, S5a, Table S6). Overall, cell morphology and variability were more associated with axial polarity and jaw placement and showed lesser location-specificity than the protein expression (Figs. 7b, S6). Among cell measures, shapes covaried most closely with axial position in both jaws, whereas polarity and size were more location-specific (Figs. 7, S6, Table S6). As condensations grew, positional and location-specific variance in protein expression diminished within the condensation, its boundary, and the overlying epithelium, but not in the uncondensed mesenchyme or its adjacent epithelium (Figs. 7a-d). Likewise, positional and location-specific variance in cell morphology declined as condensations grew, while increasing in uncondensed mesenchyme (Fig. 7d). Once condensations formed, protein expression and cell variation in their mesenchyme became uniform across axes and jaw placements (Table S6).

**Figure 7.**
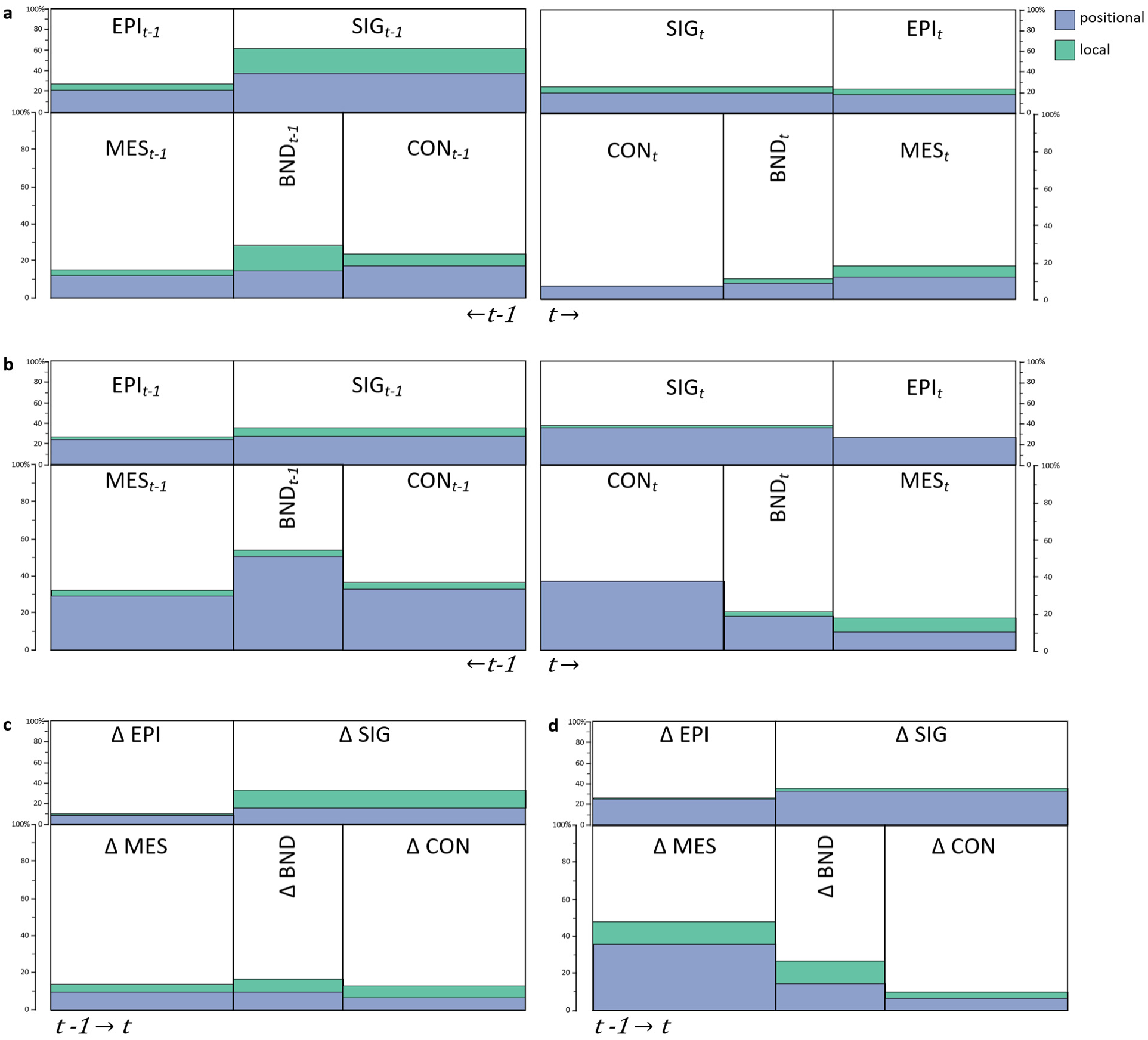
Distribution of positional (blue) and local (green) variance between pre (*t* -1) and condensation (*t*) stages in **(a)** protein expression and **(b)** cell morphology, density and variability measures (averages for cellular and molecular contributions) and changes associated with condensation formation in focal tissues in **(c)** protein expression, and **(d)** cellular measures. Data in Table S6; Figs. S5-6.

## Discussion

We found that position-specificity in avian beak mesenchymal condensations arises when some mesenchymal cells transiently match the protein expression of the progressively expanding overlying epithelium and become boundaries of future condensations (scenario Fig. 1f). Whereas signaling cross-talk between mesenchymal cells and overlying epithelium can underpin regional specificity of condensation placement (Richman and Tickle 1989; Hall and Miyake 1992; Francis-West et al. 1994; LaMantia et al. 2000; Mina et al. 2002; Haworth et al. 2004; Brito et al. 2008; Kumar et al. 2012; Lee et al. 2021; Xu et al. 2023), here we revealed the temporally resolved dynamics of this process shared by all condensations under this study. Our results further suggest that location-specificity of a condensation may be a by-product of a transient “local signal matching” phase inherent in epithelial-mesenchymal cross talk during condensation initiation. Further, our results directly implicate the mesenchymal boundary of condensations as a precise anchoring mechanism (see also Meinecke et al. 2018); the boundary could allow mesenchyme to internalize and sustain initial epithelium signaling, establishing the basis of reciprocal region-specific cross-talk between the two tissues. Finally, we find that the local signal matching phase is brief as accumulating NCM cells progressively lose their regional differences in the focal protein expression, likely restoring cell differentiation potential across region-specific condensations.

These findings raise three key questions. First, what are the mechanisms by which NCM and epithelial cells share their protein profiles and how is location-specificity in these profiles established? Second, how do accumulating NCM cells converge in their molecular profiles and erase the initial location-specificity of their tissues within condensations (Fig. 7c, d). Of particular relevance is the relationship between the number of accumulating cells and their ability to maintain their uniformity and the relationship between the condensation size, the onset of its differentiation, and the diversity of tissues it can produce. Third, what enables the conserved regulatory proteins studied here to drive context-specific and often sequentially opposite (Fig. 3a) histological changes during condensation origin and differentiation? (Fig. 3b).

Several molecular and epigenetic mechanisms could account for the observed transient signal mimicry of overlying epithelium by the adjacent mesenchyme. First, the multipotent NCM cells arriving at the sites of future condensations express a wide array of transcription factors (e.g. Dlx, Msx, Sox family genes) able to respond to epithelial cues and also possess the open chromatin architecture making epithelial-like enhancers accessible (e.g., Van Otterloo et al. 2022; Selleri and Rijli 2023). This “poised” state can enable NCM cells to rapidly match local combinations of signaling ligands under epithelial induction. Under this scenario, NCM cells receiving an epithelial signal activate a transcriptional program similar to the epithelial source which allows them to express this signal themselves (Francis-West et al. 1994; Kumar et al. 2012; Hu et al. 2015a; Xavier et al. 2016; Xu et al. 2023). This acquired autonomous signaling, in turn, either amplifies signaling through feedback to the epithelium or propagates the signal further into the adjacent mesenchyme. There are several examples of such reciprocal induction followed by signal internalization and amplification. For example, in facial mesenchyme, exposure to epithelial Shh upregulates mesenchymal hedgehog pathway components (e.g. Ptch1, Gli1) in patterns mirroring the epithelium’s Shh domain: if Shh is experimentally upregulated in NCM cells, the mesenchyme begins to behave as an epithelial anchoring source, leading to formation of extra condensations (Brito et al. 2008; Hu et al. 2015a; Xavier et al. 2016).

Second, the transient epithelium-mesenchyme matching might involve leakage of molecular signaling between the adjacent tissues or their active transport by cell rearrangements and growth (e.g., Kicheva and Briscoe 2023). For example, cells of future boundaries, that match epithelial signaling first (Fig. 4a), were more polarized towards the overlying epithelium than other mesenchymal cells (Fig. 5a). This, together with these cells’ propensity to form denser aggregations (Fig. 3b), raises the possibility that the observed epithelial signaling extension is produced by mesenchymal cellular outgrowth. Interestingly, the mesenchyme of future boundaries had a similar protein profile to the overlying epithelium, but highly distinct protein expression and polarization to its adjacent mesenchyme, including that of the future condensation (Fig. 4e), implying that it is the mesenchyme that extends the epithelial signaling and not the other way around. This directionality is strongly corroborated by the finding that protein profiles of signaling epithelium increasingly diverge from other tissues, whereas those of condensation boundary consistently converge with other tissues as condensation forms (Fig. 4e).

The finding that the signaling mimicry between adjacent tissues differs between locations (Fig. 7), raises the possibility that regional specificity of NCM cell condensations is a by-product of their reciprocal signaling interaction with local epithelium. Under this scenario, the initial anchoring of a mesenchymal condensation by epithelial mimicry is reinforced by mesenchyme’s own amplification of local signaling. A combination of molecular memory in groups of NCM cells accumulated since their induction and during migration, and developmental divergence of local epithelium produces a highly specific signaling combination; signal matching between the two adjacent tissues will thus produce location-specific condensations by default (Wu et al. 2006; Eames and Schneider 2008; Welsh and O’Brien 2009; Kaucka et al. 2016; Palmquist et al. 2022; Selleri and Rijli 2023). These mechanisms can scale up to the level of specific axial placement of condensations as long as the epithelial-mesenchymal feedback involves the induction of the boundary of a future condensation. Indeed, in the frontonasal ectodermal zone (FEZ) in avian embryos a mesenchymal signaling boundary is formed at the interface of Fgf8 and Shh domains in the overlying epithelium (Hu et al. 2003; Foppiano et al. 2007).

The underlying mesenchyme can influence and even mimic this signaling location: if mesenchymal signaling is blocked, Shh expression in the FEZ epithelium is lost. Conversely, activating Shh signaling in the mesenchyme induces an ectopic FEZ-like pattern of Shh/Fgf8 and alters condensation formation (Lu et al. 2024). Similar patterns are observed in epithelial-mesenchymal feedback of WNT/β-catenin signaling (Reid et al. 2011).

Interestingly, this location-specificity of protein profiles is short-lived and is rapidly replaced by general positional and tissue specific protein expression as condensations form. This echoes a classical finding that the initial “anchoring” matching of epithelial signaling is often brief and highly stage-specific, whereas the subsequent amplification and divergence of signaling by condensed mesenchyme, once it takes over as an autonomous signaling center, must be prolonged to maintain condensations despite rapid divergence of surrounding embryonic tissues (Hall and Miyake 1992). Essentially, the boundary of a condensation allows condensed mesenchyme to internalize the external epithelial signaling.

Distance to the boundaries also determines the rate of cellular and molecular homogenization within condensations and, correspondingly, the onset of cell differentiation once this uniformity cannot be maintained after a condensation reaches a certain size (Cottrill et al. 1987; Hall and Miyake 1992). Indeed, we found that mesenchyme cells rapidly converged in their molecular signaling and erased location-specificity in their cellular and molecular components as condensation grew (Figs. 7c,d; Table S6). The acquisition of molecular uniformity coincides with cellular transformation – condensation cells became smaller and more uniform in size, shape and polarity as they acquire similar protein expression (Figs. 4, 6, 7; Table 3, S6). Within condensations, cell uniformity is maintained through self-referencing and competition; the ability to mix and move is key to this process (Kaucka et al. 2016; Shahbazi et al. 2017; Zheng and al. 2021). Once cell movement is restricted, the condensation’s size could determine the onset and range of differentiation (Badyaev et al. 2025a). Future studies in this system can investigate the speed of reprograming of condensation cells in relation to condensation size (Angelozzi et al. 2022) as well as the fate of the boundary itself and its ability to maintain a condensation once it undergoes molecular homogenization despite acquiring distinct morphology within the condensation mesenchyme (Fig. 3b).

**Table 3.**
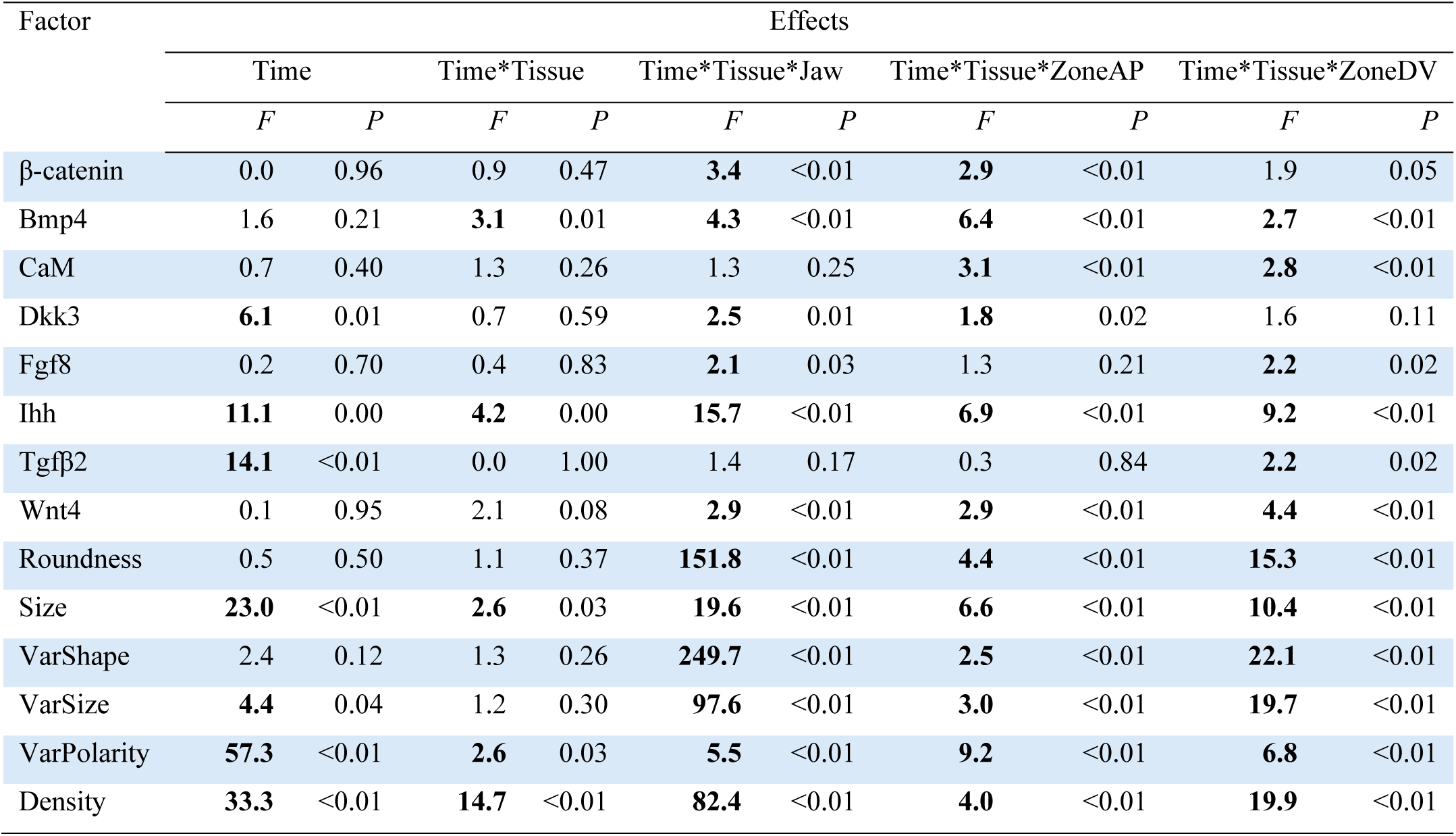
Context-dependence of change from precondensation to condensation stage in relation to tissue type, upper vs lower jaw, anterior-posterior and dorsoventral zones. Statistics as in Table 2.

| Factor | Effects |  |  |  |  |  |  |  |  |  |
| --- | --- | --- | --- | --- | --- | --- | --- | --- | --- | --- |
|  | Time |  | Time*Tissue |  | Time*Tissue*Jaw |  | Time*Tissue*ZoneAP |  | Time*Tissue*ZoneDV |  |
|  | *F* | *P* | *F* | *P* | *F* | *P* | *F* | *P* | *F* | *P* |
| β-catenin | 0.0 | 0.96 | 0.9 | 0.47 | **3.4** | <0.01 | **2.9** | <0.01 | 1.9 | 0.05 |
| Bmp4 | 1.6 | 0.21 | **3.1** | 0.01 | **4.3** | <0.01 | **6.4** | <0.01 | **2.7** | <0.01 |
| CaM | 0.7 | 0.40 | 1.3 | 0.26 | 1.3 | 0.25 | **3.1** | <0.01 | **2.8** | <0.01 |
| Dkk3 | **6.1** | 0.01 | 0.7 | 0.59 | **2.5** | 0.01 | **1.8** | 0.02 | 1.6 | 0.11 |
| Fgf8 | 0.2 | 0.70 | 0.4 | 0.83 | **2.1** | 0.03 | 1.3 | 0.21 | **2.2** | 0.02 |
| Ihh | **11.1** | 0.00 | **4.2** | 0.00 | **15.7** | <0.01 | **6.9** | <0.01 | **9.2** | <0.01 |
| Tgfb2 | **14.1** | <0.01 | 0.0 | 1.00 | 1.4 | 0.17 | 0.3 | 0.84 | **2.2** | 0.02 |
| Wnt4 | 0.1 | 0.95 | 2.1 | 0.08 | **2.9** | <0.01 | **2.9** | <0.01 | **4.4** | <0.01 |
| Roundness | 0.5 | 0.50 | 1.1 | 0.37 | **151.8** | <0.01 | **4.4** | <0.01 | **15.3** | <0.01 |
| Size | **23.0** | <0.01 | **2.6** | 0.03 | **19.6** | <0.01 | **6.6** | <0.01 | **10.4** | <0.01 |
| VarShape | 2.4 | 0.12 | 1.3 | 0.26 | **249.7** | <0.01 | **2.5** | <0.01 | **22.1** | <0.01 |
| VarSize | **4.4** | 0.04 | 1.2 | 0.30 | **97.6** | <0.01 | **3.0** | <0.01 | **19.7** | <0.01 |
| VarPolarity | **57.3** | <0.01 | **2.6** | 0.03 | **5.5** | <0.01 | **9.2** | <0.01 | **6.8** | <0.01 |
| Density | **33.3** | <0.01 | **14.7** | <0.01 | **82.4** | <0.01 | **4.0** | <0.01 | **19.9** | <0.01 |

The proteins studied here are involved in a multitude of cellular and histological processes of avian beak development (Wu et al. 2004; Betancur et al. 2010; Cheng et al. 2024), composing a core regulatory network able to produce an array of topologically plastic configurations with context-dependent effects on cellular dynamics (Mallarino et al. 2012; Badyaev et al. 2025b). In agreement with other studies, we found key involvement of Bmp4, Fgf8 and CaM in the origin of condensations (Hall and Miyake 2000; Santagati and Rijli 2003; Minoux and Rijli 2010; Svandova et al. 2020). For example, Bmp4 was upregulated at sites of future condensations (Fig. 3b, Table S4), where it promotes cell proliferation and cell compaction, whereas the strong upregulation of Fgf8 can prevent cell apoptosis at adjacent tissues in addition to providing positional information (Hall and Miyake 2000; Santagati and Rijli 2003; Wu et al. 2004; Wu et al. 2006; Barna and Niswander 2007; Hu and Marcucio 2009; Minoux and Rijli 2010). Interestingly, Bmp4 was sharply downregulated once condensations started to form (but upregulated in their boundaries, Fig. 3a), potentially to delay cell differentiation before the condensation reaches full size, as the presence of Bmp4 is associated with chondrogenesis and osteogenesis especially when coupled with high Ihh expression as found here (Fig. 3a)(Wu et al. 2006; Mallarino et al. 2012; Cela et al. 2016; Giffin et al. 2019).

Context- and time-specific modulation of the protein network can explain some discrepancies between our and previous studies. For example, a downregulation of mesenchymal Tgfβ2 was found to be required for condensation formation (Ray and Chapman 2015), whereas we found a strong upregulation of this protein immediately prior to condensation formation in the tissues of the future boundary and overlying epithelium followed by an upregulation in condensing mesenchyme (Figs. 3a, 4a; Table S4).

Although, similarly to Bmp4, downregulation of Tgfβ2 can delay the onset of cell differentiation within condensations and prevent the formation of linkages to Bmp4 and Wnt4 that promote osteogenesis (Ray and Chapman 2015), here Tgfβ2’s increase can be explained by its upregulation of CaM, which is essential for cell adhesion and compaction in the formation of condensation boundaries (Figs. 3a, 4a, Table S4) (Widelitz et al. 1993; Hall and Miyake 2000; Lin et al. 2006). Interestingly, Bmp4 and Tgfβ2 expression had the lowest positional and location-specific variance, indicating their conserved global effects compared to other focal proteins (Fig. S5). Similarly, a strong upregulation of β-catenin within future condensation boundaries was consistently associated with cell polarized aggregation and compaction (Fig. 3) whereas β-catenin increase in the overlying epithelium (Fig. 3a) suggests mechanical reorganization of epithelial tissues inducing aggregation of the adjacent mesenchymal cells (e.g., Shyer et al. 2017; Palmquist et al. 2022).

Our results also shed light on the long-range cellular processes contributing to condensation formation (scenarios *ii* and *iii* in Fig. 2a are supported by our findings). Accumulating cell density within condensations was accompanied by a relative decrease of cell density in the surrounding mesenchyme (Fig. 3b, also visible in Fig. S2) which, in combination with the strong lateral polarity of mesenchymal tissues adjacent to condensations (Fig. 5a), suggests that cell migration contributes to condensation formation and cell compaction (Fig. 2a). Further, condensation formation is associated with a relative increase in cell proliferation at the condensation site as evidenced by the decrease in average cell sizes and cell variability (Fig. 6), corroborating previous findings (Hall and Miyake 2000; Brembeck et al. 2006; Wu et al. 2006; Barna and Niswander 2007; Hall et al. 2014; Abramyan and Richman 2015; Giffin et al. 2019; Paudel et al. 2022). Cell polarity affects the orientation of cell division and cell migration and plays a key role in tissue reorganization during craniofacial development (e.g., Yamaguchi et al. 1999; Li et al. 2017). β-catenin-independent Wnt pathways affect cell polarity (Gao et al. 2011; Ho et al. 2012; Konopelski Snavely et al. 2023) and we found that distinct Wnt4 expression in the boundary, condensation, and uncondensed mesenchyme corresponded to divergent cell polarity in these tissues (Figs. 3, 4b, 5). Furthermore, the formation of condensations was associated with the opposite changes in Wnt4 expression between signaling and adjacent epithelium (Fig. 6), producing a signaling boundary between adjacent mesenchymal tissues with orthogonal polarity (Fig. 5).

In sum, the developmental organization revealed here reconciles the multipotentiality of NCM cells, evolutionarily conserved epithelial-mesenchymal signaling, and the regional specificity of cell condensations needed for proper development. Because the juxtaposition and growth dynamics of cell condensations ultimately determine evolutionary diversification in avian beak shapes and sizes (Abzhanov et al. 2004, 2006; Wu et al. 2004; Campàs et al. 2010; Badyaev 2011; Linde-Medina et al. 2016; Schneider 2024), the mechanisms underlying the onset and erasure of position-specificity in NCM stem cells are likely to be central in this process.

## Methods

### Data collection and sample sizes

We measured cell morphology and protein expression in upper and lower beak primordia of house finch embryos across developmental stages HH25-36 from five genetically distinct population groups (Table S1, Badyaev 2010; 2020). Protocols for egg collection, incubation to the required developmental stage, and field storage of samples are in (Badyaev et al. 2003; 2025a). Beaks were cryosectioned at 8 μm and stored at -80°C. Thirteen sections per individual were obtained at beak midline: one section was stained with Alcian blue hematoxylin and eosin (H&E; U. Rochester MC) to delineate the area of interest (AOI) and tissues, twelve were used in immunohistochemical (IHC) analysis of eight proteins described below.

### Tissue assignment, condensation identification, and cell morphology

Cell measurements and protein expression data were collected within AOI of the upper and lower beak that was delineated by landmarks homologous across developmental stages and confirmed with H&E histological staining (Lee et al. 2024). Briefly, the upper beak AOI was from the point of inflection of the upper beak, to the lower outer tissue of the brain/eye to the inner edge of the mouth, just past the palatine process, but not including the palatine process, to the tip of the beak, along the inner edge of the egg tooth and back to the point of upper beak inflection. The lower beak AOI was from the tip of the lower beak, down the inner edge past the developing tongue until the beak begins to widen, across the lower beak, and back to the lower beak tip along the outer edge. Four equal anterior-posterior (AP) and two equal dorsoventral (DV) zones were established within the upper and lower beaks based on the uniform size of overlaying grid (see below; Fig. 1a)

In this work, we focused only on pre-condensation and the earliest condensation stages, excluding any condensations that initiated tissue differentiation because it is associated with distinct expression of focal proteins (Ray and Chapman 2015; Cela et al. 2016). The workflow for tissue assignment and curation, and high-throughput measurements of cell morphology, variability and density, and protein expression using software we developed for this work (Lee et al. 2024) is detailed in Fig. S1. We focused on 16 mesenchymal condensations (eight in upper beak and eight in lower) that were present and repeatably identifiable in >80% of samples (Tables 1, S2, Figs. S2-4). In the upper beak, three of these condensations (U2, U5, and U8) were parts of the frontonasal facial prominence, three others (U4, U10, U12) were contained within the lateral nasal facial prominence and two (U6-7 and U9) eventually merge into the maxillary prominence. In the lower beak, six condensations will form the basihyal (L3-6) and paraglossal (L1, L5, L7) cartilages and associated muscle (L8 and L9). L2 will form the dentary bone and L4 –Meckel’s cartilage (Tables 1, S2). Within each grid box, for all cells, we measured area (μm^2^), perimeter (μm), aspect ratio 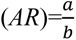 (major and minor axis of the best fitting ellipsoid), solidity, circularity, the cell shape index 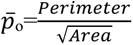, and polarity angle of the major axis (folded into 0-180° range). For each grid box (∼130 cells per box, *n* = 2,120,160 cells total), we also calculated mean and standard deviation (s.d.) for these measures. Details and repeatability of measurements are in (Badyaev et al. 2025a).

### Immunohistochemistry

We measured the expression of eight proteins that play a central role in avian beak development (Abzhanov et al. 2004, 2006; Wu et al. 2004, 2006; Mallarino et al. 2011; 2012; Fritz et al. 2014; Ray and Chapman 2015; Schneider 2018a; 2024; Badyaev et al. 2025b): β-catenin, bone morphogenic protein 4 (Bmp4), calmodulin1 (Calm1), dickkopf homolog 3 (Dkk3), fibroblast growth factor 8 (Fgf8), Indian hedgehog (Ihh), transforming growth factor beta 2 (TGFβ2), and wingless type 4 (Wnt4). For immunostaining, we used anti-β-catenin (610153, 1:16,000, BD Transduction Laboratories), anti-CaM (sc-137079, 1:15, Santa Cruz Biotechnology), anti-Wnt4 (ab91226, 1:800; Abcam), anti-Tgfβ2 (ab36495, 1:800, Abcam), anti-Bmp4 (ab118867, 1:100, Abcam), anti-Ihh (ab184624, 1:100, Abcam), anti-Dkk3 (ab214360, 1:100, Abcam), and anti-Fgf8 (89550, 1:50, Abcam) antibodies using methods described previously (Lee et al. 2024). Validations confirming specificity of stains are in (Badyaev et al. 2025a).

Reactions were visualized with either diaminobenzidine (DAB, Elite ABC HRP Kit, PK-6100, Vector Labs) or Vector Red Alkaline Phosphatase substrate and Vectastain ABC-AP Kit (AK-5000, Vector Labs) and nuclei were counterstained with hematoxylin. Three slides, each containing four tissue sections (12 sections per embryo) were run with the following grouped antibodies: i) β-catenin, Fgf8, Tgfβ2 and no primary control, ii) Bmp4, Wnt4, Ihh and no primary control, and iii) Dkk3 and no primary control, and CaM and no primary control. These were imaged and named according to embryo ID, protein, developmental stage, and IHC run to enable high-throughput processing of cell-based expression as described in (Lee et al. 2024). Protein expression was measured as the ratio of cells expressing the protein to the total number of cells within the tissue. Across IHC runs, we randomized the assignment of sections from different populations and stages.

### Statistical analyses

To achieve normal distribution, reduce skewness and stabilize variance, we used the Box-Cox transformation with λ=0.5 for raw values of protein expression, the standard logarithm transformation for cell morphology measures and the arcsine transformation for cell density and AR proportional measures. We derived linear principal components of the normalized measurements of cell morphology and variability that reduced these measurements to two dimensions – cell shape and size as well as variation in these parameters (Table S3). We used a mixed-effects model of Proc Mixed (SAS 9.4), to compute the least squares (LS) means of cell morphology and protein expression across tissue types, time periods, and condensation placements, and compared them with a Sidak adjustment for multiple comparisons.

Population and embryo identity were treated as random effects. To evaluate tissue divergence and its change between time periods, we computed pairwise Euclidean distances between the tissues in molecular (8 factors) and cellular (5 factors) divergence. We compared cell polarity angles with Watson-Williams axial test and adjusted multiple pairwise tissue comparisons with Bonferroni correction.

In addition to the strong positional effects of placement along AP and DV axes across jaws or tissues, the majority of cellular and molecular mechanisms also showed substantial context-dependency as evidenced by significant two- and three-way interactions among these predictors (see below). This gave us an opportunity to directly compare the contribution of positional effects (zones along AP and DV axes and jaw placement, Fig. 1a), local, or region-specific “local” effects (interactions between these predictors) and of effects not accounted for by either gradient or local placement (which includes tissue-specificity of cellular and molecular mechanisms; Fig. 2). For every tissue type, we compared the explanatory power of context-specific (“local”) effects in cellular and molecular responses by contrasting the marginal R² from a full model – which included all predictors and a full array of their interactions – with that of a reduced model which excluded all interaction terms. Both models were estimated using maximum likelihood approach in mixed-effects models, in which fixed effects captured the primary predictors with Proc PLM(SAS 9.4). Fixed-effect predictions were generated and their variance computed, while random and residual variances were extracted from the covariance parameters. We calculated the marginal R² as the ratio of the variance due to fixed effects to the total variance (the sum of fixed, random, and residual variances). Contribution of the context-specific “local” predictors was the difference of marginal R²s between the full and reduced models, contribution of positional effects was contribution of fixed positional (non-interaction positional predictors, i.e., zones and jaws) to the full model, and global “unexplained” variance was calculated as the variance in cellular or regulatory effect, not explained by the model (Fig. 2b,c).

## Data availability

All data are available in the manuscript and the supplementary materials.

## Acknowledgments

We thank C. Lee, S. Britton, F. Bravo, K. Gahl, K. Chenard, C. Seliga, X. Posner, R. Hollingsworth, G. Semenov, J. Abtahi, O. Puebla, and A. Shaikh for help with collection and preparation of samples, cryosectioning, histological and molecular assays and computational work.

## Supplementary Materials

**Figure S1.**
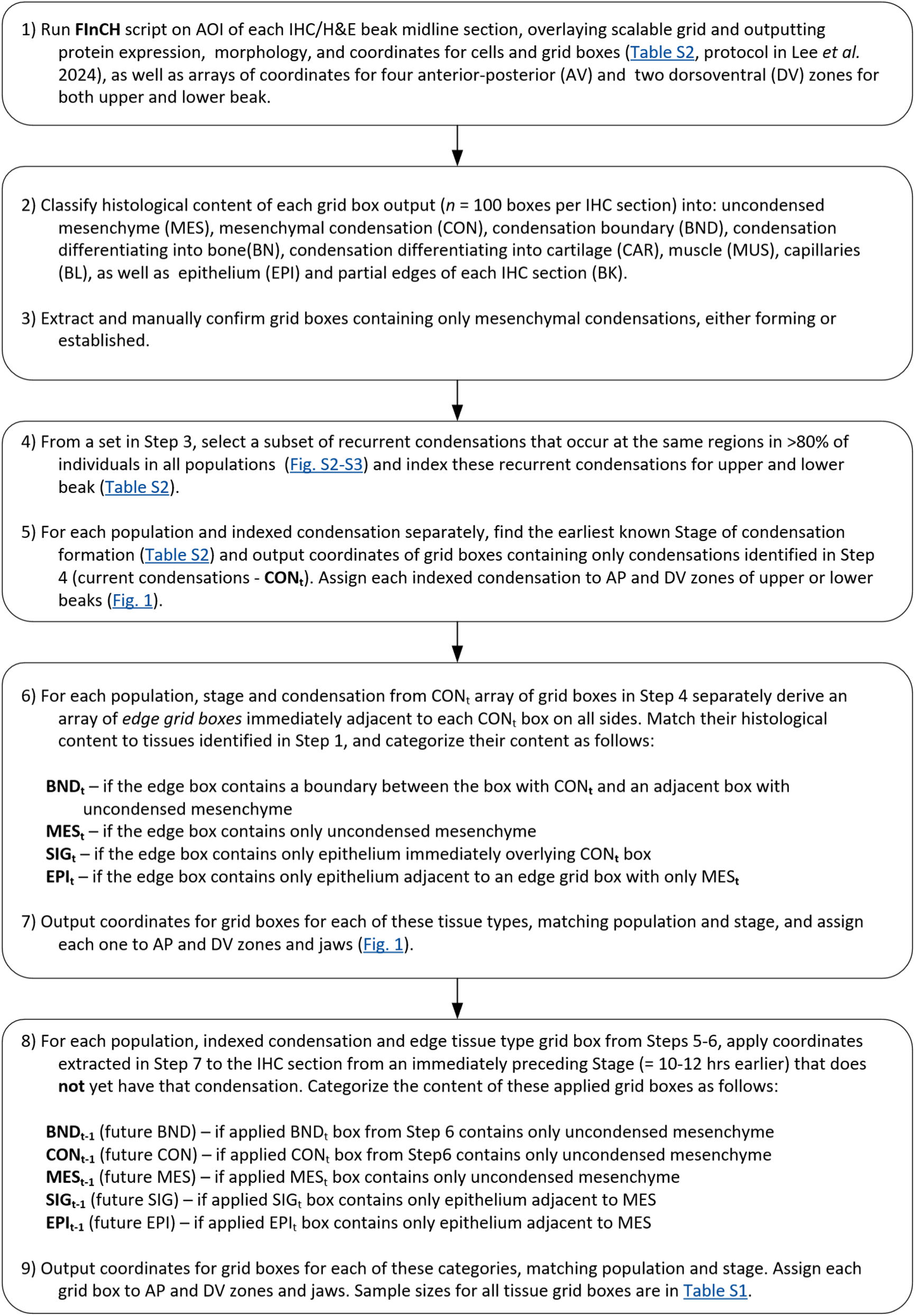
Workflow for tissue assignment for pre- (*t*-1) and condensation (*t*) stages for Fig 1a insert.

**Figure S2.**
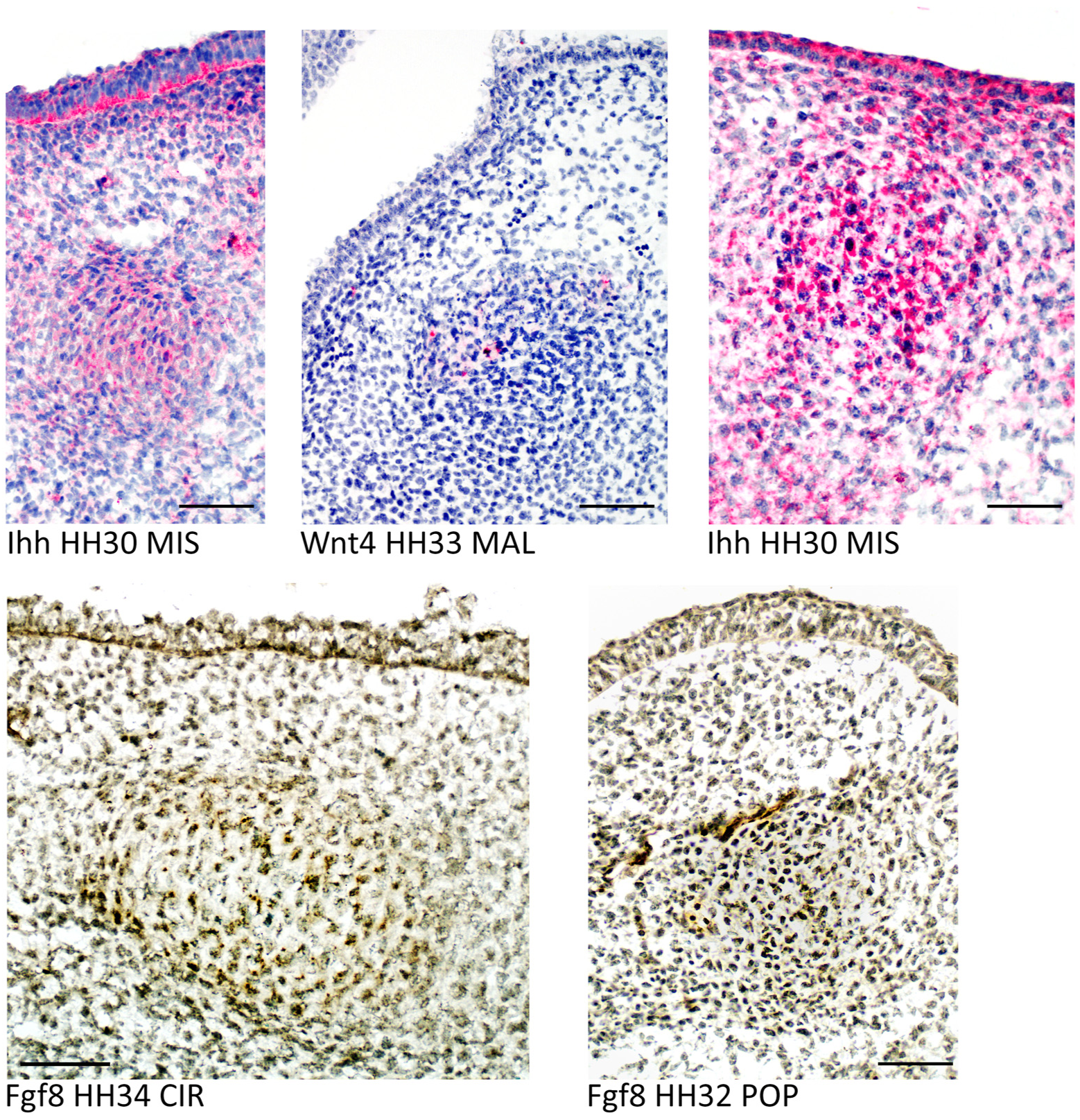
Examples of mesenchymal condensations and overlying epithelium investigated in this study. Text shows protein expression, developmental stage and population abbreviation. Scale bar is 50 μm.

**Figure S3.**
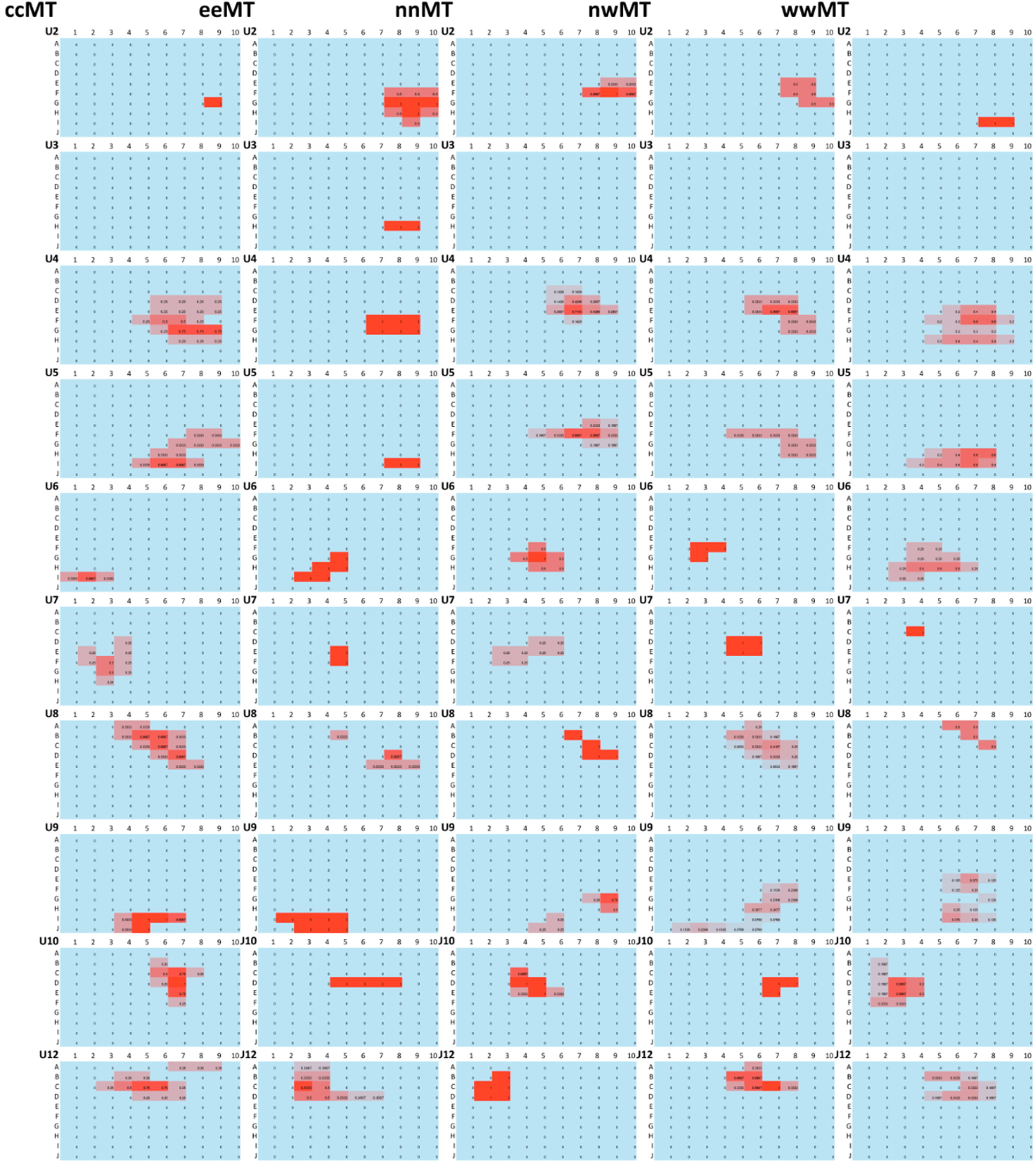
Frequency of occurrence for the upper beak condensations across populations. Shown are the locations of the earliest occurrence. Intensity of red color and numbers indicates proportion of samples with condensation location in this grid box. Summary in Table S2.

**Figure S4.**
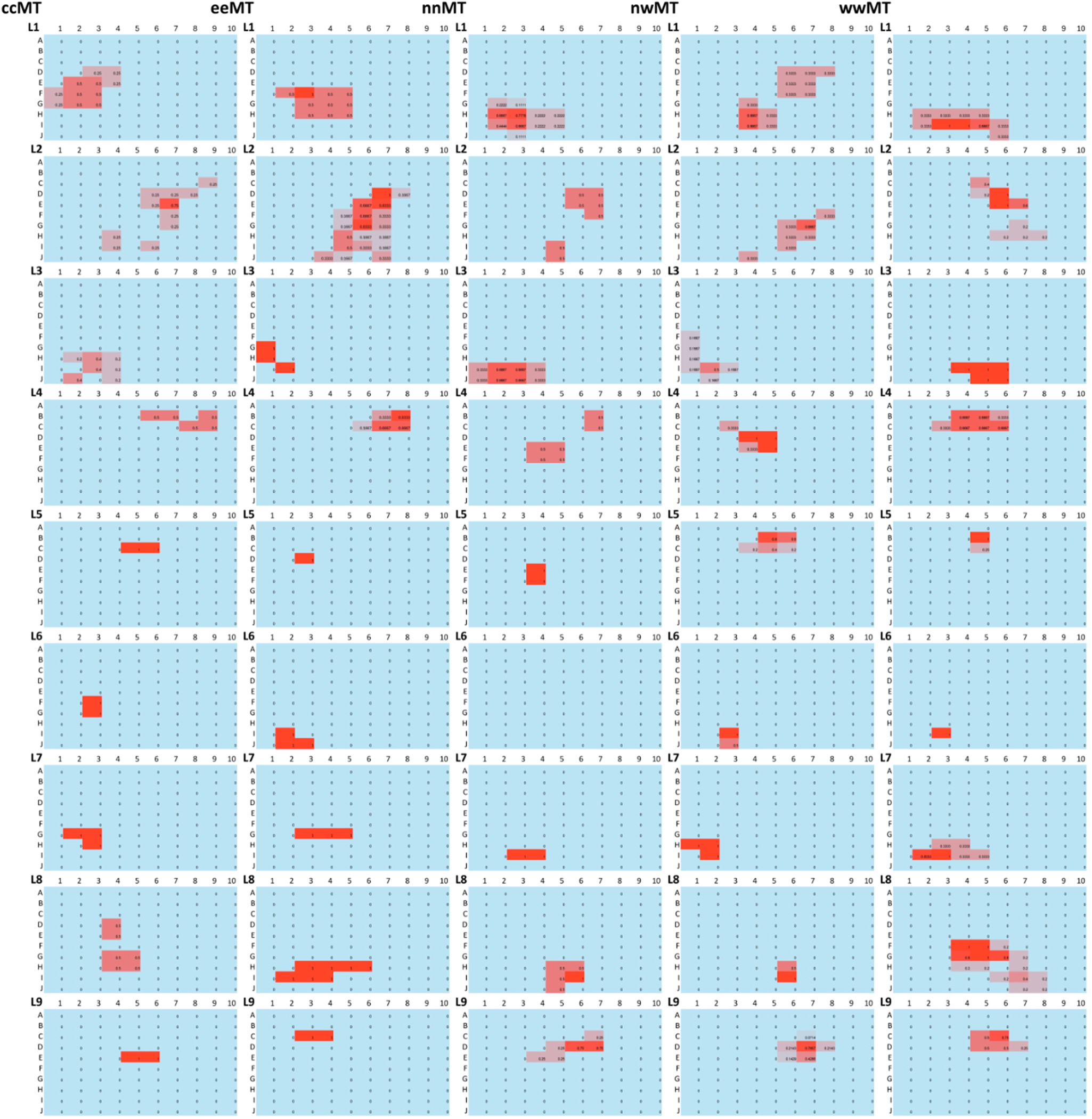
Frequency of occurrence for the lower beak condensations across populations. Shown are the locations of the earliest occurrence. Intensity of red color and numbers indicates proportion of samples with condensation location in this grid box. Summary in Table S2.

**Figure S5.**
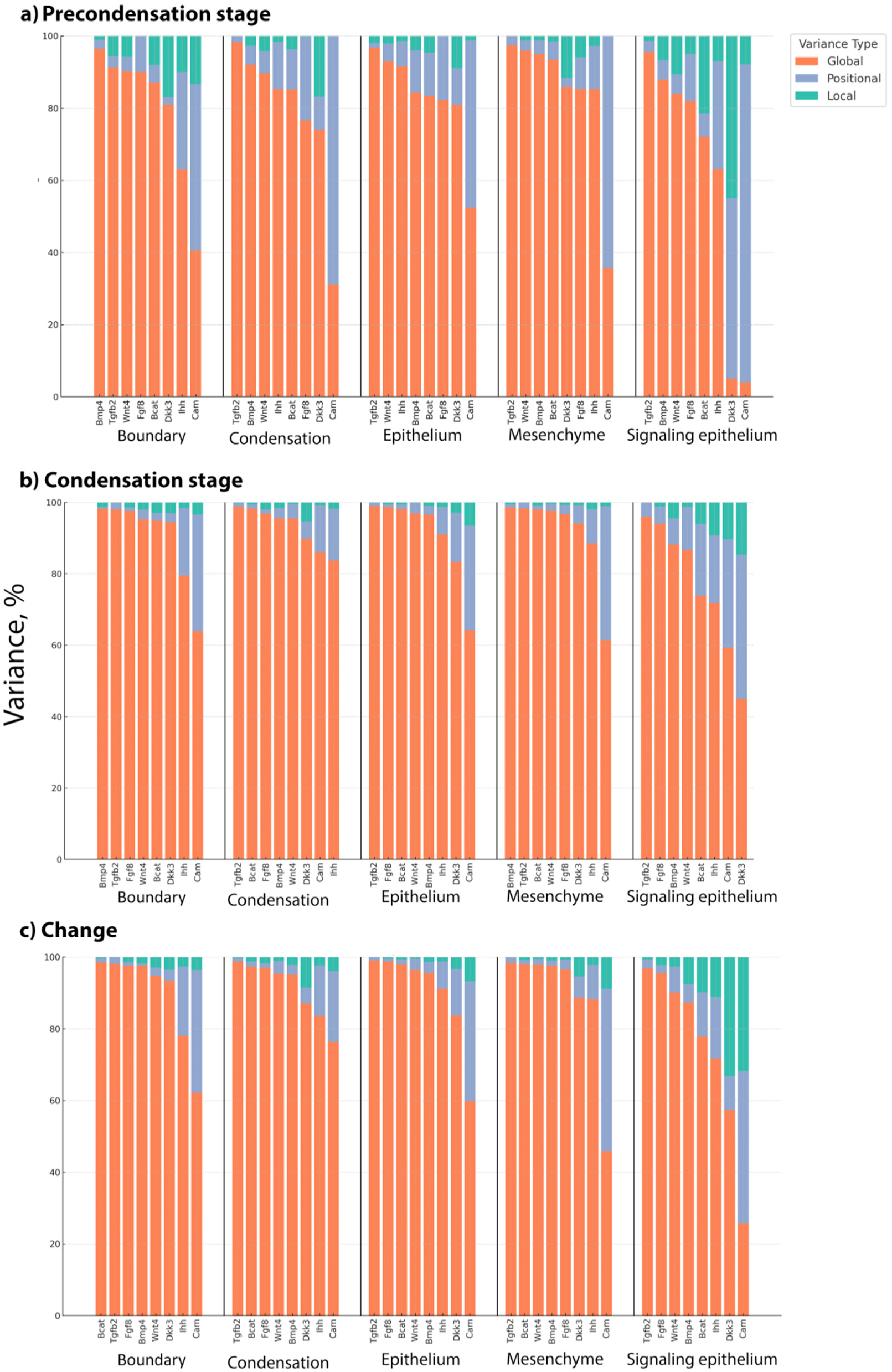
Partitioning of variance in protein expression across tissues due to global, positional, and local variance contribution during **(a)** precondensation stage, (**b**) condensation stage and (**c**) in change from pre- to condensation stage. Within each tissue, proteins are arranged by declining “global” variance. Within-tissue averages are shown in Fig. 7, Table S6.

**Figure S6.**
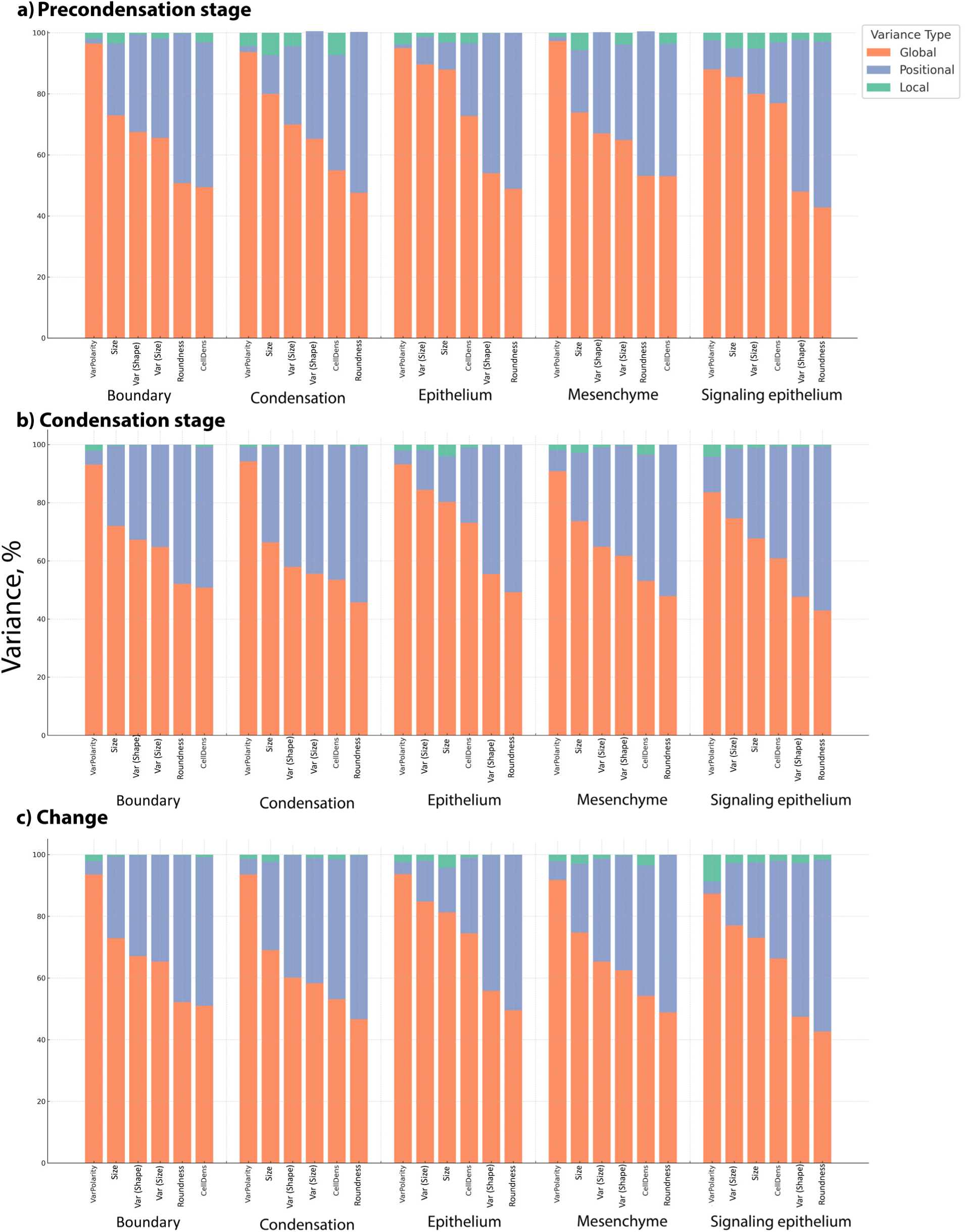
Partitioning of variance in cell morphology and variability across tissues between global, positional, and local variance sources during **(a)** precondensation stage, (**b**) condensation stage and (**c**) in change from pre- to condensation stage. Within each tissue, cell measures are arranged by declining “global” variance. Within-tissue averages are shown in Fig. 7, Table S6.

**Table S1.** Sample sizes of embryo samples and condensation grid boxes per developmental stage.

| Region<br>**abbreviation**<br>*total embryos* | Sampling locations<br>latitude, °N, longitude, °W)<br>within a region | Developmental stages (HH)<br>*grid boxes* |  |  |  |
| --- | --- | --- | --- | --- | --- |
|  |  | 25-29 | 30-32 | 33-34 | 35-36 |
| Central Montana<br>**cMT**<br>*n* = 17 | Fairfield (47.615545, 111.98016)<br>Great Falls (47.49839, 111.22444)<br>Cut Bank (48.629812, 112.333318)<br>Choteau (47.81437, 112.19193)<br>Valier (48.305658, 112.251628)<br>Lima (44.6369°, 112.5920°) | 1,128 | 2,750 | 1,232 | 1,507 |
| Eastern Montana<br>**eMT**<br>*n* = 35 | Plentywood (48.77368, 104.547787)<br>Glasgow (48.202811, 106.634599)<br>Circle (47.4156, 105.5907)<br>Culbertson (48.147585, 104.514955)<br>Poplar (48.1131°, 105.1983°)<br>Wolf Point (48.0906°, 105.6406°) | 1,009 | 6,708 | 3,324 | 1,105 |
| Northern Montana<br>**nMT**<br>*n* = 19 | Havre (48.5500, 109.6841)<br>Malta (48.352662, 107.880582) | 1,513 | 2,516 | 1,933 | 194 |
| Northwest Montana<br>**nwMT**<br>*n* = 25 | Kalispell (48.21943, 114.33305)<br>Polson (47.687447, 114.160314)<br>Superior (47.193045, 114.888817)<br>Plains (47.459956, 114.884694)<br>Thompson Falls (47.600191, 115.342068) | 339 | 3,607 | 2,654 | 819 |
| Western Montana<br>**wMT**<br>*n* = 10 | Missoula (46.8787, 113.9966)<br>Paws Up (46.916957, 113.435576)<br>Hamilton (46.246863, 114.173587) | 563 | 1,336 | 1,518 | 0 |
| Total:<br>*n* = 106 embryos (*n* = 35,755 grid box samples) |  | 4,552 | 16,917 | 10,661 | 3,625 |

**Table S2.** Centroids (X, Y, and centroid grid cell) of condensations at earliest and maximum appearances with corresponding stages.

| CON | Region | Early |  |  |  | Maximum |  |  |  | CON | Region | Early |  |  |  | Maximum |  |  |  |
| --- | --- | --- | --- | --- | --- | --- | --- | --- | --- | --- | --- | --- | --- | --- | --- | --- | --- | --- | --- |
|  |  | Centroid |  |  | Cell | Centroid |  |  | Cell |  |  | Centroid |  |  | Cell | Centroid |  |  | Cell |
|  |  | Stage | X | Y |  | Stage | X | Y |  |  |  | Stage | X | Y |  | Stage | X | Y |  |
| L1 | ccMT | 25 | 5.8 | 2.5 | F3 | 35 | 6.3 | 3.5 | F4 | U10 | ccMT | 31 | 3.8 | 6.8 | D7 | 35 | 5.0 | 3.9 | E4 |
| L1 | eeMT | 27 | 6.8 | 3.7 | G4 | 35 | 6.9 | 2.3 | G2 | U10 | eeMT | 28 | 4.0 | 6.5 | D6 | 35 | 4.0 | 4.3 | D4 |
| L1 | nnMT | 29 | 8.4 | 3.0 | H3 | 35 | 8.1 | 3.3 | H3 | U10 | nnMT | 29 | 4.2 | 4.6 | D5 | 35 | 4.2 | 4.6 | D5 |
| L1 | nwMT | 30 | 6.6 | 5.5 | G5 | 36 | 6.6 | 5.5 | G5 | U10 | nwMT | 30 | 4.3 | 7.3 | D7 | 36 | 4.4 | 3.7 | D4 |
| L1 | wwMT | 27 | 8.8 | 3.9 | I4 | 34 | 5.8 | 2.7 | F3 | U10 | wwMT | 32 | 4.6 | 3.0 | E3 | 34 | 4.2 | 3.1 | D3 |
| L2 | ccMT | 25 | 5.7 | 6.5 | F7 | 35 | 5.8 | 5.3 | F5 | U12 | ccMT | 29 | 2.7 | 5.7 | C6 | 35 | 2.7 | 5.7 | C6 |
| L2 | eeMT | 28 | 6.7 | 6.2 | G6 | 35 | 6.7 | 6.2 | G6 | U12 | eeMT | 28 | 3.1 | 3.8 | C4 | 35 | 2.8 | 4.8 | C5 |
| L2 | nnMT | 26 | 6.1 | 6.1 | F6 | 35 | 7.0 | 5.4 | G5 | U12 | nnMT | 29 | 3.2 | 2.6 | C3 | 35 | 4.1 | 4.7 | D5 |
| L2 | nwMT | 30 | 7.7 | 6.4 | H6 | 36 | 6.6 | 6.8 | G7 | U12 | nwMT | 30 | 2.5 | 6.2 | B6 | 36 | 3.2 | 2.4 | C2 |
| L2 | wwMT | 27 | 5.1 | 6.2 | E6 | 34 | 6.1 | 7.1 | F7 | U12 | wwMT | 30 | 3.1 | 6.4 | C6 | 33 | 3.5 | 1.4 | D1 |
| L3 | ccMT | 31 | 8.9 | 3.0 | I3 | 35 | 8.9 | 3.0 | I3 | U2 | ccMT | 29 | 7.0 | 9.0 | G9 | 35 | 6.6 | 9.1 | G9 |
| L3 | eeMT | 30 | 8.0 | 1.3 | H1 | 35 | 8.1 | 1.4 | H1 | U2 | eeMT | 27 | 7.2 | 9.0 | G9 | 35 | 7.2 | 9.0 | G9 |
| L3 | nnMT | 31 | 9.5 | 2.5 | J2 | 33 | 9.5 | 2.5 | J2 | U2 | nnMT | 26 | 5.8 | 9.1 | F9 | 35 | 6.8 | 9.5 | G9 |
| L3 | nwMT | 31 | 8.4 | 1.7 | H2 | 36 | 8.4 | 2.0 | H2 | U2 | nwMT | 26 | 6.0 | 8.8 | F9 | 36 | 6.0 | 8.8 | F9 |
| L3 | wwMT | 27 | 9.4 | 5.2 | I5 | 34 | 9.4 | 5.2 | I5 | U2 | wwMT | 27 | 9.0 | 8.5 | I8 | 34 | 7.6 | 9.3 | H9 |
| L4 | ccMT | 25 | 2.4 | 7.8 | B8 | 35 | 2.7 | 7.9 | C8 | U3 | eeMT | 33 | 8.0 | 8.5 | H8 | 33 | 8.0 | 8.5 | H8 |
| L4 | eeMT | 28 | 2.6 | 7.5 | C7 | 35 | 2.5 | 7.7 | B8 | U4 | ccMT | 25 | 6.1 | 7.4 | F7 | 35 | 4.9 | 5.6 | E6 |
| L4 | nnMT | 26 | 4.5 | 5.3 | D5 | 33 | 3.7 | 6.4 | D6 | U4 | eeMT | 27 | 6.5 | 8.0 | F8 | 35 | 6.5 | 8.0 | F8 |
| L4 | nwMT | 26 | 4.3 | 4.5 | D4 | 36 | 4.5 | 7.5 | D8 | U4 | nnMT | 29 | 4.6 | 7.2 | E7 | 35 | 4.7 | 5.5 | E6 |
| L4 | wwMT | 27 | 2.6 | 4.8 | C5 | 33 | 2.6 | 4.8 | C5 | U4 | nwMT | 30 | 5.2 | 7.6 | E8 | 36 | 5.0 | 5.7 | E6 |
| L5 | ccMT | 25 | 3.0 | 5.5 | C6 | 35 | 3.9 | 4.0 | D4 | U4 | wwMT | 27 | 6.6 | 7.1 | G7 | 34 | 4.5 | 5.2 | D5 |
| L5 | eeMT | 28 | 4.0 | 3.0 | D3 | 35 | 3.5 | 4.3 | D4 | U5 | ccMT | 25 | 7.9 | 7.4 | H7 | 35 | 5.6 | 4.8 | F5 |
| L5 | nnMT | 31 | 5.5 | 4.0 | F4 | 33 | 4.1 | 3.8 | D4 | U5 | eeMT | 27 | 9.0 | 8.5 | I8 | 35 | 6.4 | 7.1 | F7 |
| L5 | nwMT | 31 | 2.4 | 5.3 | B5 | 36 | 4.6 | 3.9 | E4 | U5 | nnMT | 29 | 5.9 | 7.6 | F8 | 35 | 6.7 | 7.2 | G7 |
| L5 | wwMT | 32 | 2.2 | 5.0 | B5 | 34 | 5.1 | 4.0 | E4 | U5 | nwMT | 30 | 6.8 | 7.5 | G8 | 36 | 6.9 | 5.7 | G6 |
| L6 | ccMT | 31 | 6.5 | 3.0 | F3 | 35 | 6.5 | 3.0 | F3 | U5 | wwMT | 27 | 8.5 | 6.6 | I7 | 34 | 8.5 | 6.6 | I7 |
| L6 | eeMT | 32 | 9.7 | 2.3 | J2 | 34 | 9.7 | 2.3 | J2 | U6 | ccMT | 33 | 9.0 | 2.0 | I2 | 35 | 8.8 | 2.7 | I3 |
| L6 | nwMT | 34 | 9.3 | 3.0 | I3 | 35 | 8.1 | 3.7 | H4 | U6 | eeMT | 30 | 8.2 | 4.2 | H4 | 34 | 8.2 | 4.2 | H4 |
| L6 | wwMT | 34 | 9.0 | 3.0 | I3 | 34 | 9.0 | 3.0 | I3 | U6 | nnMT | 29 | 7.1 | 5.1 | G5 | 33 | 7.1 | 5.1 | G5 |
| L7 | ccMT | 25 | 7.3 | 2.7 | G3 | 35 | 8.2 | 2.6 | H3 | U6 | nwMT | 31 | 6.3 | 3.3 | F3 | 36 | 5.6 | 4.2 | F4 |
| L7 | eeMT | 27 | 7.0 | 4.0 | G4 | 35 | 8.5 | 3.5 | H4 | U6 | wwMT | 30 | 7.7 | 4.7 | H5 | 30 | 7.7 | 4.7 | H5 |
| L7 | nnMT | 31 | 9.0 | 3.5 | I4 | 33 | 9.0 | 3.1 | I3 | U7 | ccMT | 31 | 6.1 | 3.2 | F3 | 35 | 6.1 | 3.2 | F3 |
| L7 | nwMT | 30 | 8.3 | 1.7 | H2 | 36 | 7.8 | 2.5 | H2 | U7 | eeMT | 30 | 5.5 | 5.0 | F5 | 35 | 6.8 | 3.8 | G4 |
| L7 | wwMT | 30 | 8.8 | 3.2 | I3 | 34 | 8.8 | 3.2 | I3 | U7 | nnMT | 29 | 5.0 | 4.5 | E4 | 33 | 6.0 | 4.5 | F4 |
| L8 | ccMT | 25 | 6.5 | 4.3 | F4 | 35 | 7.6 | 3.5 | H3 | U7 | nwMT | 30 | 4.5 | 5.5 | D6 | 36 | 4.5 | 5.5 | D6 |
| L8 | eeMT | 27 | 8.4 | 3.9 | H4 | 35 | 5.6 | 5.1 | F5 | U7 | wwMT | 30 | 3.0 | 4.0 | C4 | 33 | 5.5 | 3.3 | F3 |
| L8 | nnMT | 26 | 8.8 | 5.5 | I6 | 33 | 7.5 | 3.4 | G3 | U8 | ccMT | 32 | 2.8 | 5.9 | C6 | 35 | 3.4 | 5.6 | C6 |
| L8 | nwMT | 26 | 8.7 | 6.0 | I6 | 36 | 7.1 | 3.8 | G4 | U8 | eeMT | 28 | 4.2 | 7.5 | D8 | 35 | 2.5 | 5.0 | B5 |
| L8 | wwMT | 27 | 7.2 | 5.4 | G5 | 34 | 7.2 | 5.4 | G5 | U8 | nnMT | 26 | 3.3 | 8.0 | C8 | 35 | 3.0 | 5.5 | C6 |
| L9 | ccMT | 25 | 5.0 | 5.5 | E6 | 35 | 3.5 | 6.2 | C6 | U8 | nwMT | 31 | 3.0 | 6.6 | C7 | 36 | 2.3 | 4.7 | B5 |
| L9 | eeMT | 28 | 3.0 | 3.5 | C4 | 35 | 5.0 | 6.3 | E6 | U8 | wwMT | 30 | 1.8 | 7.0 | B7 | 34 | 2.5 | 3.5 | B4 |
| L9 | nnMT | 31 | 4.1 | 6.0 | D6 | 33 | 4.1 | 6.0 | D6 | U9 | ccMT | 29 | 9.3 | 5.4 | I5 | 35 | 9.3 | 5.4 | I5 |
| L9 | nwMT | 31 | 4.3 | 6.9 | D7 | 36 | 4.6 | 5.2 | E5 | U9 | eeMT | 28 | 9.4 | 3.7 | I4 | 35 | 9.4 | 3.7 | I4 |
| L9 | wwMT | 30 | 3.5 | 5.7 | D6 | 34 | 3.5 | 5.7 | D6 | U9 | nnMT | 29 | 8.1 | 7.8 | H8 | 35 | 9.1 | 4.3 | I4 |
|  |  |  |  |  |  |  |  |  |  | U9 | nwMT | 31 | 8.1 | 6.0 | H6 | 36 | 8.1 | 6.0 | H6 |
|  |  |  |  |  |  |  |  |  |  | U9 | wwMT | 30 | 7.1 | 6.8 | G7 | 33 | 7.1 | 6.8 | G7 |

**Table S3.** Principal components of six measures of cell morphology and variability (correlational matrix) within *n* = 35,755 grid boxes (*n* = 2.12 x 10^6^ cells, ∼130 cells/grid box) containing forming condensation. Bold values show the largest projection on the original variable. Prin1s mostly captures cell elongation (left) and its variability (right) and Prin2s – cell size (left) and its variability (right).

| Variable | Prin 1<br>*roundness* | Prin 2<br>*size* | Variable | Prin 1<br>*roundness (var)* | Prin 2<br>*size (var)* |
| --- | --- | --- | --- | --- | --- |
| Area_Mean | 0.29 | **0.68** | Area_SD | -0.18 | **0.73** |
| Shape_Mean | **-0.51** | 0.09 | Shape_SD | **0.51** | 0.01 |
| Aspect Ratio_Mean | **-0.44** | -0.09 | Aspect Ratio_SD | **0.42** | 0.12 |
| Perimeter_Mean | -0.17 | **0.71** | Perimeter_SD | 0.27 | **0.66** |
| Circularity_Mean | **0.50** | -0.09 | Circularity_SD | **0.48** | -0.10 |
| Solidity_Mean | **0.51** | 0.05 | Solidity_SD | **0.48** | -0.11 |
| Eigenvalue | 3.73 | 1.73 | Eigenvalue | 3.72 | 1.58 |
| Variance explained | 62.3% | 29.8% | Variance explained | 62.0% | 26.4% |

**Table S4.**
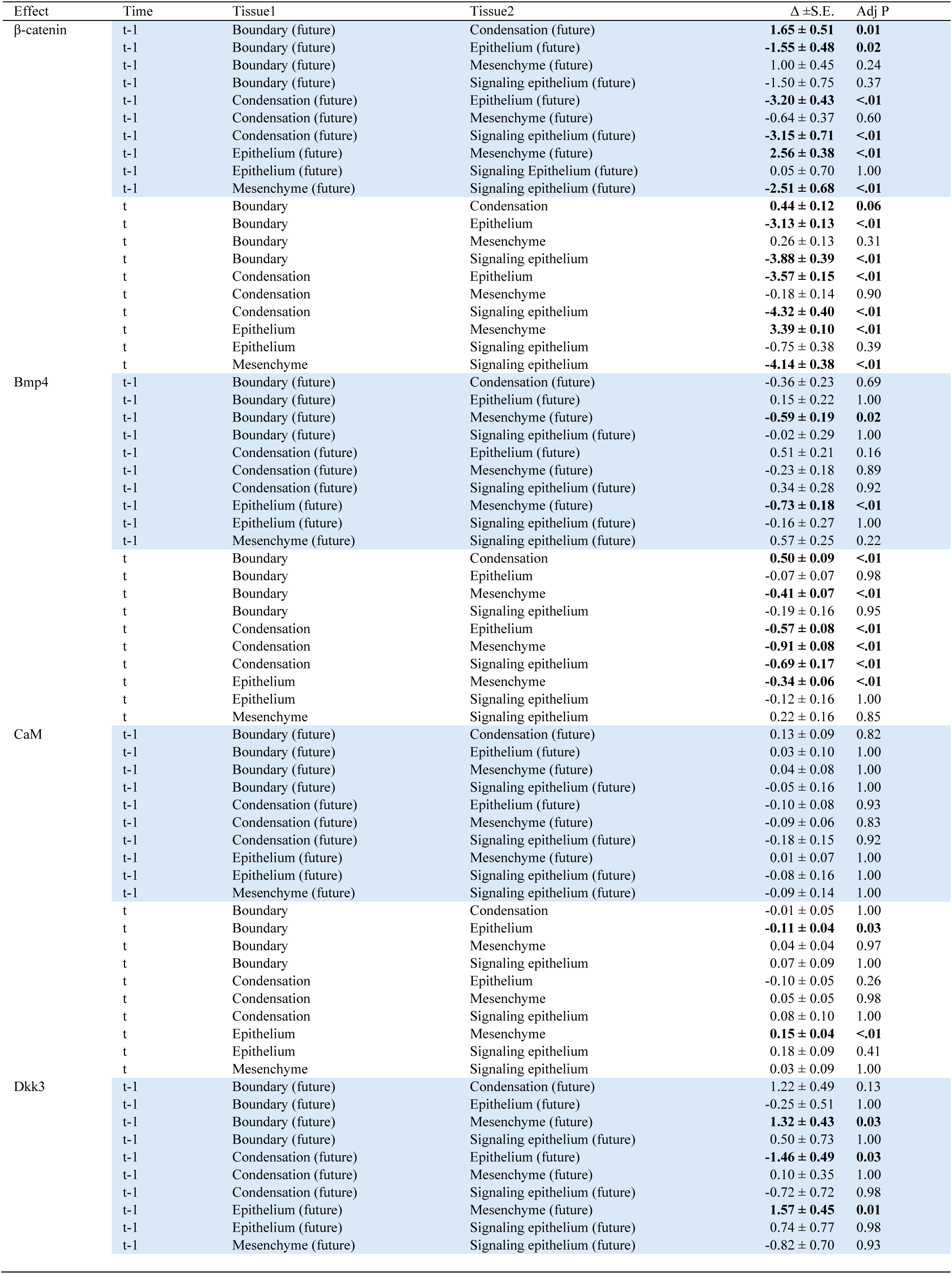
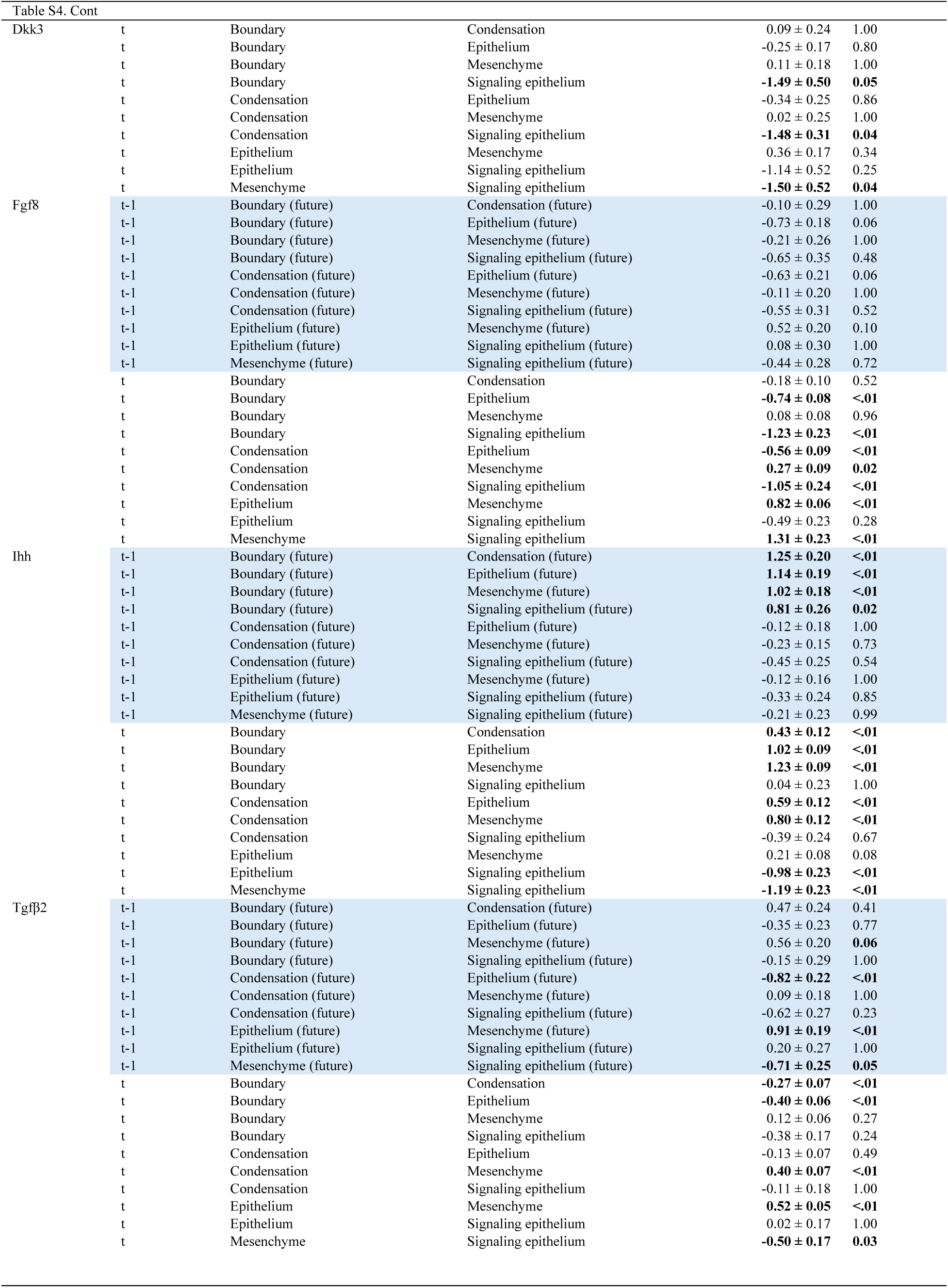
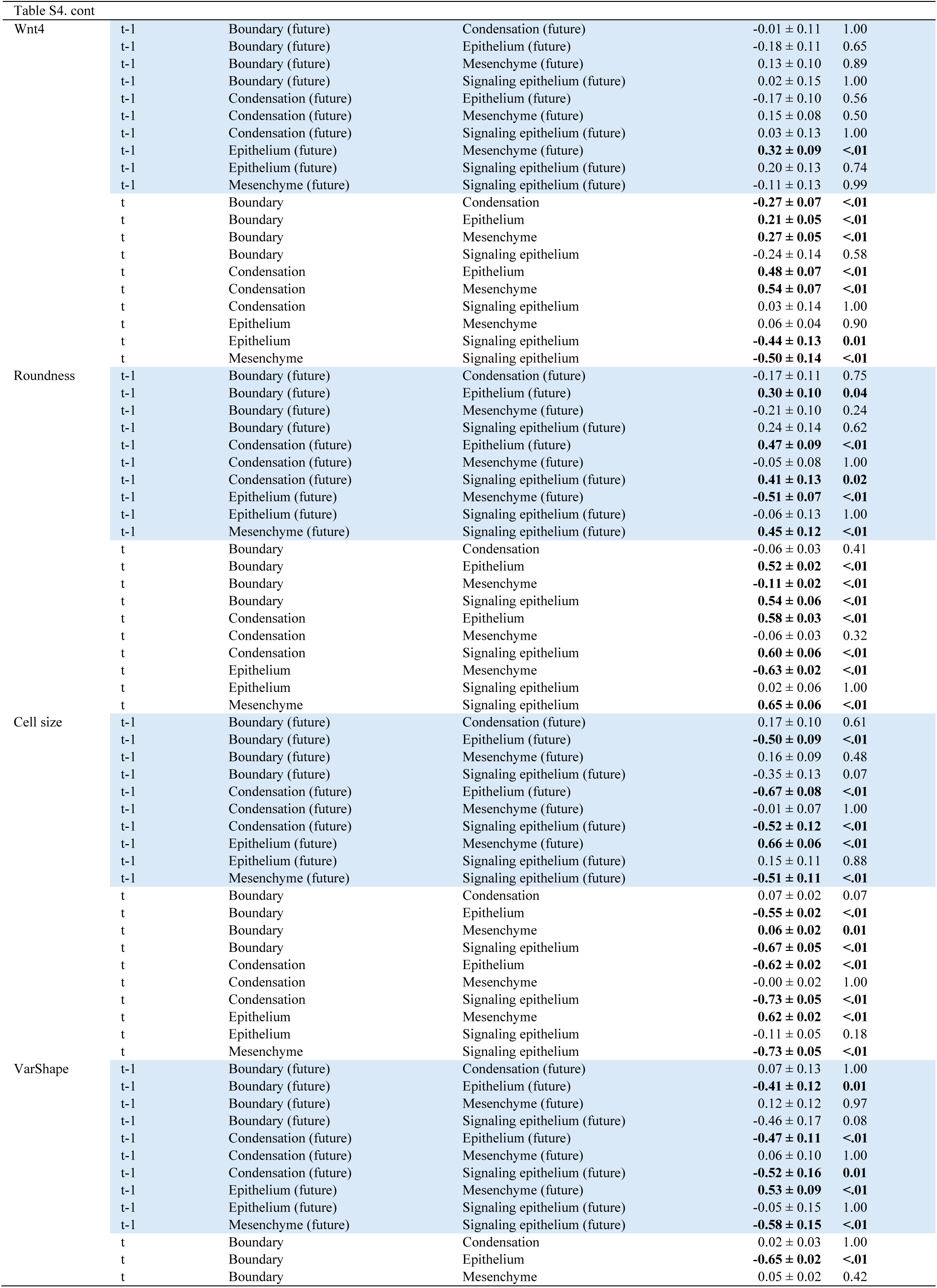
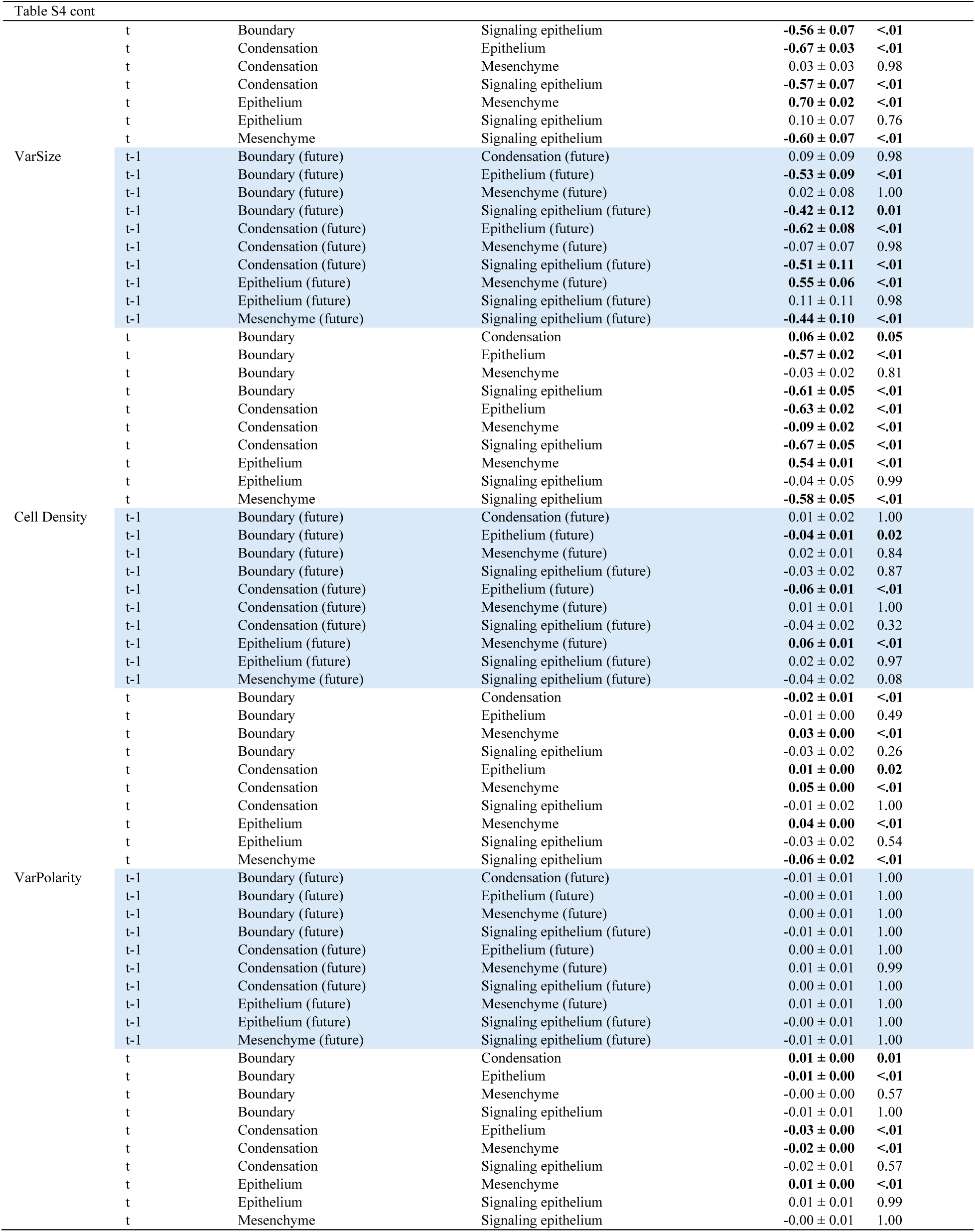
Differences of least squares means (controlling for the effects of Population, Jaw, ZoneAP, Zone DV and embryo ID) for protein expression and cell morphology immediately prior (future) and during (present) condensation formation.

**Table S5.**
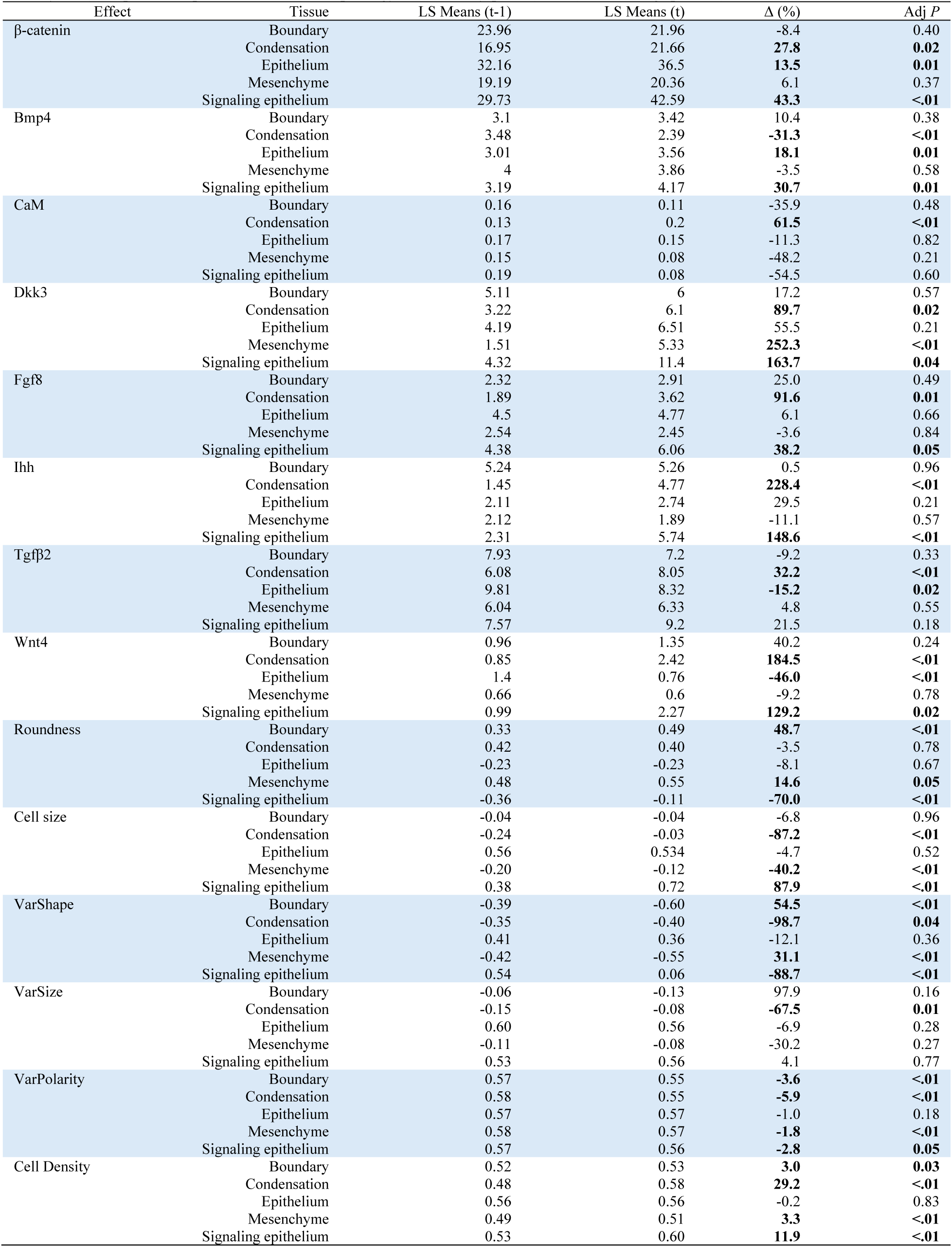
Relative difference (%) in least squares means (Controlling for the effects of Population, Jaw, ZoneAP, Zone DV and embryo ID) for protein expression and cell morphology immediately prior (future) and during (present) condensation formation.

| Effect | Tissue | LS Means (t-1) | LS Means (t) | $\Delta$ (%) | Adj *P* |
| --- | --- | --- | --- | --- | --- |
| β-catenin | Boundary | 23.96 | 21.96 | -8.4 | 0.40 |
|  | Condensation | 16.95 | 21.66 | **27.8** | **0.02** |
|  | Epithelium | 32.16 | 36.5 | **13.5** | **0.01** |
|  | Mesenchyme | 19.19 | 20.36 | 6.1 | 0.37 |
|  | Signaling epithelium | 29.73 | 42.59 | **43.3** | **<.01** |
| Bmp4 | Boundary | 3.1 | 3.42 | 10.4 | 0.38 |
|  | Condensation | 3.48 | 2.39 | **-31.3** | **<.01** |
|  | Epithelium | 3.01 | 3.56 | **18.1** | **0.01** |
|  | Mesenchyme | 4 | 3.86 | -3.5 | 0.58 |
|  | Signaling epithelium | 3.19 | 4.17 | **30.7** | **0.01** |
| CaM | Boundary | 0.16 | 0.11 | -35.9 | 0.48 |
|  | Condensation | 0.13 | 0.2 | **61.5** | **<.01** |
|  | Epithelium | 0.17 | 0.15 | -11.3 | 0.82 |
|  | Mesenchyme | 0.15 | 0.08 | -48.2 | 0.21 |
|  | Signaling epithelium | 0.19 | 0.08 | -54.5 | 0.60 |
| Dkk3 | Boundary | 5.11 | 6 | 17.2 | 0.57 |
|  | Condensation | 3.22 | 6.1 | **89.7** | **0.02** |
|  | Epithelium | 4.19 | 6.51 | 55.5 | 0.21 |
|  | Mesenchyme | 1.51 | 5.33 | **252.3** | **<.01** |
|  | Signaling epithelium | 4.32 | 11.4 | **163.7** | **0.04** |
| Fgf8 | Boundary | 2.32 | 2.91 | 25.0 | 0.49 |
|  | Condensation | 1.89 | 3.62 | **91.6** | **0.01** |
|  | Epithelium | 4.5 | 4.77 | 6.1 | 0.66 |
|  | Mesenchyme | 2.54 | 2.45 | -3.6 | 0.84 |
|  | Signaling epithelium | 4.38 | 6.06 | **38.2** | **0.05** |
| Ihh | Boundary | 5.24 | 5.26 | 0.5 | 0.96 |
|  | Condensation | 1.45 | 4.77 | **228.4** | **<.01** |
|  | Epithelium | 2.11 | 2.74 | 29.5 | 0.21 |
|  | Mesenchyme | 2.12 | 1.89 | -11.1 | 0.57 |
|  | Signaling epithelium | 2.31 | 5.74 | **148.6** | **<.01** |
| Tgfb2 | Boundary | 7.93 | 7.2 | -9.2 | 0.33 |
|  | Condensation | 6.08 | 8.05 | **32.2** | **<.01** |
|  | Epithelium | 9.81 | 8.32 | **-15.2** | **0.02** |
|  | Mesenchyme | 6.04 | 6.33 | 4.8 | 0.55 |
|  | Signaling epithelium | 7.57 | 9.2 | 21.5 | 0.18 |
| Wnt4 | Boundary | 0.96 | 1.35 | 40.2 | 0.24 |
|  | Condensation | 0.85 | 2.42 | **184.5** | **<.01** |
|  | Epithelium | 1.4 | 0.76 | **-46.0** | **<.01** |
|  | Mesenchyme | 0.66 | 0.6 | -9.2 | 0.78 |
|  | Signaling epithelium | 0.99 | 2.27 | **129.2** | **0.02** |
| Roundness | Boundary | 0.33 | 0.49 | **48.7** | **<.01** |
|  | Condensation | 0.42 | 0.40 | -3.5 | 0.78 |
|  | Epithelium | -0.23 | -0.23 | -8.1 | 0.67 |
|  | Mesenchyme | 0.48 | 0.55 | **14.6** | **0.05** |
|  | Signaling epithelium | -0.36 | -0.11 | **-70.0** | **<.01** |
| Cell size | Boundary | -0.04 | -0.04 | -6.8 | 0.96 |
|  | Condensation | -0.24 | -0.03 | **-87.2** | **<.01** |
|  | Epithelium | 0.56 | 0.534 | -4.7 | 0.52 |
|  | Mesenchyme | -0.20 | -0.12 | **-40.2** | **<.01** |
|  | Signaling epithelium | 0.38 | 0.72 | **87.9** | **<.01** |
| VarShape | Boundary | -0.39 | -0.60 | **54.5** | **<.01** |
|  | Condensation | -0.35 | -0.40 | **-98.7** | **0.04** |
|  | Epithelium | 0.41 | 0.36 | -12.1 | 0.36 |
|  | Mesenchyme | -0.42 | -0.55 | **31.1** | **<.01** |
|  | Signaling epithelium | 0.54 | 0.06 | **-88.7** | **<.01** |
| VarSize | Boundary | -0.06 | -0.13 | 97.9 | 0.16 |
|  | Condensation | -0.15 | -0.08 | **-67.5** | **0.01** |
|  | Epithelium | 0.60 | 0.56 | -6.9 | 0.28 |
|  | Mesenchyme | -0.11 | -0.08 | -30.2 | 0.27 |
|  | Signaling epithelium | 0.53 | 0.56 | 4.1 | 0.77 |
| VarPolarity | Boundary | 0.57 | 0.55 | **-3.6** | **<.01** |
|  | Condensation | 0.58 | 0.55 | **-5.9** | **<.01** |
|  | Epithelium | 0.57 | 0.57 | -1.0 | 0.18 |
|  | Mesenchyme | 0.58 | 0.57 | **-1.8** | **<.01** |
|  | Signaling epithelium | 0.57 | 0.56 | **-2.8** | **0.05** |
| Cell Density | Boundary | 0.52 | 0.53 | **3.0** | **0.03** |
|  | Condensation | 0.48 | 0.58 | **29.2** | **<.01** |
|  | Epithelium | 0.56 | 0.56 | -0.2 | 0.83 |
|  | Mesenchyme | 0.49 | 0.51 | **3.3** | **<.01** |
|  | Signaling epithelium | 0.53 | 0.60 | **11.9** | **<.01** |

**Table S6.** Variance components for cellular and molecular processes associated with condensation formation. Shown are R2 of full model and percent of variance contributed by Local (2 and 3 order interactions between positional terms), Positional (gradient, axial, and jaw placement) and Global (unexplained by either positional or interaction terms) variation. AVGCELL and AVGMOL – are averages of cellular and molecular processes, correspondingly, contribution for each tissue.

| Tissue | Variable | Full R2 | Global | Positional | Local |
| --- | --- | --- | --- | --- | --- |
| Boundary (future) | Shape1 | 0.493 | 50.71 | 49.05 | 0.24 |
|  | Shape2 | 0.271 | 72.94 | 23.47 | 3.59 |
|  | ShapeUni1 | 0.325 | 67.54 | 31.82 | 0.64 |
|  | ShapeUni2 | 0.344 | 65.57 | 32.51 | 1.92 |
|  | VarPolarity | 0.036 | 96.43 | 1.50 | 2.07 |
|  | CellDens | 0.506 | 49.42 | 47.49 | 3.10 |
|  | AVGCELL | 0.329 | 67.10 | 30.97 | 1.93 |
| Condensation (future) | Shape1 | 0.525 | 47.54 | 52.74 | -0.28 |
|  | Shape2 | 0.200 | 80.00 | 12.61 | 7.40 |
|  | ShapeUni1 | 0.348 | 65.25 | 35.28 | -0.52 |
|  | ShapeUni2 | 0.301 | 69.93 | 25.74 | 4.33 |
|  | VarPolarity | 0.063 | 93.67 | 1.98 | 4.35 |
|  | CellDens | 0.451 | 54.93 | 37.75 | 7.32 |
|  | AVGCELL | 0.314 | 68.55 | 27.68 | 3.77 |
| Epithelium (future) | Shape1 | 0.512 | 48.82 | 51.13 | 0.05 |
|  | Shape2 | 0.121 | 87.93 | 8.92 | 3.15 |
|  | ShapeUni1 | 0.460 | 53.97 | 45.83 | 0.20 |
|  | ShapeUni2 | 0.104 | 89.60 | 9.00 | 1.40 |
|  | VarPolarity | 0.050 | 95.03 | 1.11 | 3.86 |
|  | CellDens | 0.272 | 72.77 | 23.69 | 3.54 |
|  | AVGCELL | 0.253 | 74.69 | 23.28 | 2.03 |
| Mesenchyme (future) | Shape1 | 0.469 | 53.14 | 47.31 | -0.45 |
|  | Shape2 | 0.261 | 73.90 | 20.42 | 5.68 |
|  | ShapeUni1 | 0.329 | 67.08 | 33.11 | -0.19 |
|  | ShapeUni2 | 0.351 | 64.86 | 31.19 | 3.94 |
|  | VarPolarity | 0.028 | 97.24 | 1.35 | 1.41 |
|  | CellDens | 0.470 | 53.04 | 43.27 | 3.68 |
|  | AVGCELL | 0.318 | 68.21 | 29.44 | 2.35 |
| Signaling epithelium (future) | Shape1 | 0.573 | 42.72 | 54.35 | 2.93 |
|  | Shape2 | 0.146 | 85.44 | 9.55 | 5.00 |
|  | ShapeUni1 | 0.520 | 47.99 | 49.66 | 2.35 |
|  | ShapeUni2 | 0.200 | 79.99 | 14.82 | 5.20 |
|  | VarPolarity | 0.119 | 88.05 | 9.42 | 2.53 |
|  | CellDens | 0.231 | 76.88 | 19.92 | 3.19 |
|  | AVGCELL | 0.298 | 70.18 | 26.29 | 3.53 |
| Boundary | Shape1 | 0.479 | 52.11 | 47.67 | 0.22 |
|  | Shape2 | 0.280 | 72.00 | 27.65 | 0.35 |
|  | ShapeUni1 | 0.328 | 67.21 | 32.52 | 0.26 |
|  | ShapeUni2 | 0.352 | 64.75 | 35.20 | 0.05 |
|  | VarPolarity | 0.069 | 93.08 | 4.69 | 2.23 |
|  | CellDens | 0.492 | 50.83 | 48.43 | 0.75 |
|  | AVGCELL | 0.333 | 66.66 | 32.69 | 0.64 |
| Condensation | Shape1 | 0.542 | 45.77 | 53.73 | 0.50 |
|  | Shape2 | 0.336 | 66.37 | 33.03 | 0.61 |
|  | ShapeUni1 | 0.421 | 57.92 | 41.94 | 0.14 |
|  | ShapeUni2 | 0.445 | 55.53 | 44.25 | 0.22 |
|  | VarPolarity | 0.058 | 94.18 | 4.97 | 0.85 |
|  | CellDens | 0.465 | 53.47 | 46.26 | 0.27 |
|  | AVGCELL | 0.378 | 62.20 | 37.36 | 0.43 |
| Epithelium | Shape1 | 0.508 | 49.16 | 50.74 | 0.10 |
|  | Shape2 | 0.197 | 80.29 | 15.70 | 4.02 |
|  | ShapeUni1 | 0.446 | 55.43 | 44.35 | 0.22 |
|  | ShapeUni2 | 0.156 | 84.37 | 13.68 | 1.95 |
|  | VarPolarity | 0.068 | 93.19 | 4.73 | 2.09 |
|  | CellDens | 0.269 | 73.13 | 25.80 | 1.07 |
|  | AVGCELL | 0.274 | 72.59 | 25.83 | 1.58 |
| Mesenchyme | Shape1 | 0.521 | 47.87 | 52.07 | 0.06 |
|  | Shape2 | 0.264 | 73.64 | 23.57 | 2.79 |
|  | ShapeUni1 | 0.383 | 61.69 | 37.94 | 0.37 |
|  | ShapeUni2 | 0.351 | 64.86 | 34.10 | 1.03 |
|  | VarPolarity | 0.092 | 90.76 | 7.29 | 1.95 |
|  | CellDens | 0.469 | 53.09 | 43.49 | 3.42 |
|  | AVGCELL | 0.347 | 65.32 | 33.08 | 1.60 |

| Table S6 cont. |  |  |  |  |  |
| --- | --- | --- | --- | --- | --- |
| Signaling epithelium | Shape1 | 0.571 | 42.91 | 56.52 | 0.57 |
|  | Shape2 | 0.323 | 67.66 | 31.15 | 1.20 |
|  | ShapeUni1 | 0.524 | 47.59 | 51.67 | 0.74 |
|  | ShapeUni2 | 0.254 | 74.61 | 24.02 | 1.38 |
|  | VarPolarity | 0.165 | 83.53 | 12.28 | 4.19 |
|  | CellDens | 0.392 | 60.81 | 38.59 | 0.60 |
|  | AVGCELL | 0.371 | 62.85 | 35.70 | 1.45 |
| Boundary (change) | Shape1 | 0.47817 | 52.18 | 47.59 | 0.23 |
|  | Shape2 | 0.27127 | 72.87 | 26.58 | 0.54 |
|  | ShapeUni1 | 0.32947 | 67.05 | 32.70 | 0.25 |
|  | ShapeUni2 | 0.34703 | 65.30 | 34.54 | 0.16 |
|  | VarPolarity | 0.064755 | 93.52 | 4.32 | 2.16 |
|  | CellDens | 0.49074 | 50.93 | 48.22 | 0.86 |
|  | AVGCELL | 0.330 | 66.98 | 32.32 | 0.70 |
| Condensation (change) | Shape1 | 0.53379 | 46.62 | 53.08 | 0.29 |
|  | Shape2 | 0.31041 | 68.96 | 28.69 | 2.35 |
|  | ShapeUni1 | 0.39862 | 60.14 | 39.66 | 0.20 |
|  | ShapeUni2 | 0.41743 | 58.26 | 40.69 | 1.05 |
|  | VarPolarity | 0.064549 | 93.55 | 5.11 | 1.34 |
|  | CellDens | 0.46896 | 53.10 | 45.35 | 1.54 |
|  | AVGCELL | 0.366 | 63.44 | 35.43 | 1.13 |
| Epithelium (change) | Shape1 | 0.50516 | 49.48 | 50.41 | 0.11 |
|  | Shape2 | 0.18708 | 81.29 | 14.44 | 4.27 |
|  | ShapeUni1 | 0.44184 | 55.82 | 43.97 | 0.21 |
|  | ShapeUni2 | 0.15276 | 84.72 | 13.24 | 2.04 |
|  | VarPolarity | 0.063001 | 93.70 | 3.83 | 2.47 |
|  | CellDens | 0.25531 | 74.47 | 24.51 | 1.03 |
|  | AVGCELL | 0.268 | 73.25 | 25.07 | 1.69 |
| Mesenchyme (change) | Shape1 | 0.51192 | 48.81 | 51.13 | 0.06 |
|  | Shape2 | 0.25265 | 74.74 | 22.23 | 3.04 |
|  | ShapeUni1 | 0.37583 | 62.42 | 37.24 | 0.34 |
|  | ShapeUni2 | 0.34686 | 65.31 | 33.41 | 1.28 |
|  | VarPolarity | 0.082946 | 91.71 | 6.14 | 2.16 |
|  | CellDens | 0.45797 | 54.20 | 42.23 | 3.57 |
|  | AVGCELL | 0.338 | 66.20 | 32.06 | 1.74 |
| Signaling epithelium (change) | Shape1 | 0.57426 | 42.57 | 55.62 | 1.81 |
|  | Shape2 | 0.26927 | 73.07 | 24.31 | 2.62 |
|  | ShapeUni1 | 0.52555 | 47.45 | 49.76 | 2.80 |
|  | ShapeUni2 | 0.22982 | 77.02 | 20.20 | 2.78 |
|  | VarPolarity | 0.12665 | 87.34 | 3.88 | 8.79 |
|  | CellDens | 0.3371 | 66.29 | 31.66 | 2.05 |
|  | AVGCELL | 0.344 | 65.62 | 30.90 | 3.47 |
| Boundary (future) | Bcat | 0.130 | 87.00 | 4.95 | 8.50 |
|  | Bmp4 | 0.036 | 96.42 | 2.52 | 1.06 |
|  | Cam | 0.594 | 40.61 | 46.03 | 13.36 |
|  | Dkk3 | 0.190 | 81.00 | 2.05 | 17.00 |
|  | Fgf8 | 0.100 | 90.00 | 10.00 | 0.00 |
|  | Ihh | 0.370 | 63.00 | 27.00 | 10.00 |
|  | Tgfb2 | 0.087 | 91.30 | 3.10 | 6.00 |
|  | Wnt4 | 0.098 | 90.20 | 4.00 | 6.00 |
|  | AVGMOL | 0.201 | 79.94 | 12.46 | 7.74 |
| Condensation (future) | Bcat | 0.149 | 85.11 | 11.14 | 3.75 |
|  | Bmp4 | 0.079 | 92.08 | 5.16 | 2.77 |
|  | Cam | 0.689 | 31.13 | 68.88 | -0.02 |
|  | Dkk3 | 0.260 | 74.02 | 9.12 | 16.86 |
|  | Fgf8 | 0.234 | 76.65 | 23.21 | 0.14 |
|  | Ihh | 0.147 | 85.26 | 13.10 | 1.64 |
|  | Tgfb2 | 0.017 | 98.32 | 1.94 | -0.26 |
|  | Wnt4 | 0.103 | 89.70 | 6.07 | 4.23 |
|  | AVGMOL | 0.210 | 79.03 | 17.33 | 3.64 |
| Epithelium (future) | Bcat | 0.166 | 83.36 | 12.02 | 4.63 |
|  | Bmp4 | 0.157 | 84.28 | 11.71 | 4.01 |
|  | Cam | 0.476 | 52.45 | 46.31 | 1.25 |
|  | Dkk3 | 0.191 | 80.93 | 10.17 | 8.90 |
|  | Fgf8 | 0.177 | 82.25 | 17.75 | 0.00 |
|  | Ihh | 0.085 | 91.54 | 6.98 | 1.48 |
|  | Tgfb2 | 0.032 | 96.77 | 1.31 | 1.92 |
|  | Wnt4 | 0.070 | 93.02 | 4.85 | 2.13 |
|  | AVGMOL | 0.169 | 83.07 | 13.89 | 3.04 |
| Mesenchyme (future) | Bcat | 0.066 | 93.45 | 5.02 | 1.54 |
|  | Bmp4 | 0.050 | 94.96 | 3.79 | 1.25 |
|  | Cam | 0.633 | 35.71 | 64.37 | 0.00 |
|  | Dkk3 | 0.142 | 85.77 | 2.62 | 11.61 |
|  | Fgf8 | 0.147 | 85.31 | 8.75 | 5.94 |
|  | Ihh | 0.147 | 85.27 | 11.94 | 2.78 |
|  | Tgfb2 | 0.025 | 97.46 | 2.53 | 0.01 |
|  | Wnt4 | 0.041 | 95.95 | 2.81 | 1.25 |
|  | AVGMOL | 0.156 | 84.36 | 12.73 | 2.91 |
| Signaling epithelium (future) | Bcat | 0.279 | 72.09 | 6.50 | 22.00 |
|  | Bmp4 | 0.121 | 87.91 | 5.36 | 6.73 |
|  | Cam | 0.960 | 4.00 | 88.14 | 8.00 |
|  | Dkk3 | 0.950 | 5.02 | 50.00 | 45.00 |
|  | Fgf8 | 0.180 | 82.00 | 13.00 | 5.00 |
|  | Ihh | 0.370 | 63.00 | 30.00 | 7.00 |
|  | Tgfb2 | 0.045 | 95.51 | 3.02 | 1.47 |
|  | Wnt4 | 0.160 | 84.00 | 5.40 | 11.00 |
|  | AVGMOL | 0.383 | 61.69 | 25.18 | 13.27 |
| Boundary | Bcat | 0.016 | 95.00 | 2.00 | 3.00 |
|  | Bmp4 | 0.017 | 98.27 | 0.48 | 1.24 |
|  | Cam | 0.360 | 63.95 | 32.69 | 3.36 |
|  | Dkk3 | 0.055 | 94.52 | 2.43 | 3.05 |
|  | Fgf8 | 0.024 | 97.58 | 0.98 | 1.45 |
|  | Ihh | 0.205 | 79.49 | 18.94 | 1.57 |
|  | Tgfb2 | 0.020 | 98.04 | 1.85 | 0.11 |
|  | Wnt4 | 0.047 | 95.26 | 2.69 | 2.04 |
|  | AVGMOL | 0.093 | 90.69 | 7.70 | 1.73 |
| Condensation | Bcat | 0.017 | 98.28 | 1.19 | 0.53 |
|  | Bmp4 | 0.044 | 95.62 | 2.84 | 1.54 |
|  | Cam | 0.140 | 86.04 | 13.17 | 0.79 |
|  | Dkk3 | 0.102 | 89.81 | 4.84 | 5.35 |
|  | Fgf8 | 0.032 | 96.85 | 1.19 | 1.96 |
|  | Ihh | 0.163 | 83.72 | 14.51 | 1.77 |
|  | Tgfb2 | 0.012 | 98.85 | 0.95 | 0.20 |
|  | Wnt4 | 0.045 | 95.52 | 4.09 | 0.40 |
|  | AVGMOL | 0.069 | 93.09 | 5.35 | 1.57 |
| Epithelium | Bcat | 0.018 | 98.17 | 1.26 | 0.57 |
|  | Bmp4 | 0.034 | 96.63 | 2.39 | 0.97 |
|  | Cam | 0.358 | 64.23 | 29.28 | 6.49 |
|  | Dkk3 | 0.166 | 83.39 | 13.65 | 2.96 |
|  | Fgf8 | 0.013 | 98.69 | 0.82 | 0.49 |
|  | Ihh | 0.089 | 91.09 | 7.64 | 1.27 |
|  | Tgfb2 | 0.010 | 99.05 | 0.90 | 0.05 |
|  | Wnt4 | 0.031 | 96.93 | 2.91 | 0.16 |
|  | AVGMOL | 0.090 | 91.02 | 7.36 | 1.62 |
| Mesenchyme | Bcat | 0.020 | 98.00 | 1.19 | 0.81 |
|  | Bmp4 | 0.014 | 98.62 | 0.84 | 0.54 |
|  | Cam | 0.386 | 61.44 | 37.49 | 1.07 |
|  | Dkk3 | 0.058 | 94.15 | 5.08 | 0.77 |
|  | Fgf8 | 0.033 | 96.72 | 2.62 | 0.66 |
|  | Ihh | 0.115 | 88.51 | 9.63 | 1.86 |
|  | Tgfb2 | 0.017 | 98.29 | 1.69 | 0.02 |
|  | Wnt4 | 0.024 | 97.65 | 1.98 | 0.37 |
|  | AVGMOL | 0.083 | 91.67 | 7.57 | 0.76 |
| Signaling epithelium | Bcat | 0.261 | 73.88 | 20.12 | 6.01 |
|  | Bmp4 | 0.118 | 88.23 | 7.34 | 4.43 |
|  | Cam | 0.407 | 59.27 | 30.47 | 10.26 |
|  | Dkk3 | 0.550 | 45.00 | 40.34 | 15.00 |
|  | Fgf8 | 0.060 | 94.04 | 4.78 | 1.18 |
|  | Ihh | 0.281 | 71.91 | 18.88 | 9.22 |
|  | Tgfb2 | 0.029 | 97.11 | 3.11 | -0.22 |
|  | Wnt4 | 0.133 | 86.67 | 12.06 | 1.27 |
|  | AVGMOL | 0.230 | 77.01 | 17.14 | 5.89 |
| Boundary (change) | Bcat | 0.015 | 98.50 | 1.09 | 0.41 |
|  | Bmp4 | 0.024 | 97.57 | 0.59 | 1.84 |
|  | Cam | 0.378 | 62.19 | 34.15 | 3.66 |
|  | Dkk3 | 0.066 | 93.37 | 3.07 | 3.56 |
|  | Fgf8 | 0.024 | 97.58 | 1.00 | 1.43 |
|  | Ihh | 0.220 | 77.99 | 19.30 | 2.72 |
|  | Tgfb2 | 0.020 | 98.01 | 1.78 | 0.20 |
|  | Wnt4 | 0.052 | 94.76 | 2.26 | 2.99 |
|  | AVGMOL | 0.100 | 89.99 | 7.91 | 2.10 |
| Condensation (change) | Bcat | 0.028 | 97.24 | 1.51 | 1.25 |
|  | Bmp4 | 0.049 | 95.06 | 2.67 | 2.27 |
|  | Cam | 0.236 | 76.36 | 19.69 | 3.94 |
|  | Dkk3 | 0.129 | 87.09 | 4.35 | 8.56 |
|  | Fgf8 | 0.030 | 97.00 | 1.27 | 1.73 |
|  | Ihh | 0.164 | 83.59 | 14.04 | 2.37 |
|  | Tgfb2 | 0.013 | 98.71 | 0.98 | 0.31 |
|  | Wnt4 | 0.046 | 95.38 | 3.48 | 1.14 |
|  | AVGMOL | 0.087 | 91.30 | 6.00 | 2.70 |
| Epithelium (change) | Bcat | 0.021 | 97.89 | 1.43 | 0.68 |
|  | Bmp4 | 0.045 | 95.49 | 3.10 | 1.41 |
|  | Cam | 0.402 | 59.78 | 33.48 | 6.74 |
|  | Dkk3 | 0.163 | 83.70 | 12.85 | 3.45 |
|  | Fgf8 | 0.013 | 98.69 | 0.85 | 0.46 |
|  | Ihh | 0.089 | 91.10 | 7.60 | 1.30 |
|  | Tgfb2 | 0.010 | 99.01 | 0.85 | 0.14 |
|  | Wnt4 | 0.036 | 96.38 | 3.14 | 0.48 |
|  | AVGMOL | 0.097 | 90.26 | 7.91 | 1.83 |
| Mesenchyme (change) | Bcat | 0.021 | 97.94 | 1.19 | 0.87 |
|  | Bmp4 | 0.024 | 97.57 | 1.39 | 1.04 |
|  | Cam | 0.541 | 45.90 | 45.22 | 8.88 |
|  | Dkk3 | 0.113 | 88.65 | 5.89 | 5.46 |
|  | Fgf8 | 0.035 | 96.49 | 2.74 | 0.76 |
|  | Ihh | 0.118 | 88.22 | 9.51 | 2.27 |
|  | Tgfb2 | 0.017 | 98.34 | 1.62 | 0.04 |
|  | Wnt4 | 0.022 | 97.77 | 1.67 | 0.57 |
|  | AVGMOL | 0.111 | 88.86 | 8.65 | 2.49 |
| Signaling epithelium (change) | Bcat | 0.222 | 77.79 | 12.35 | 9.86 |
|  | Bmp4 | 0.127 | 87.30 | 5.10 | 7.60 |
|  | Cam | 0.742 | 25.83 | 42.38 | 31.79 |
|  | Dkk3 | 0.426 | 57.36 | 9.45 | 33.20 |
|  | Fgf8 | 0.045 | 95.47 | 2.23 | 2.30 |
|  | Ihh | 0.283 | 71.72 | 17.06 | 11.22 |
|  | Tgfb2 | 0.032 | 96.84 | 2.48 | 0.68 |
|  | Wnt4 | 0.099 | 90.11 | 7.17 | 2.72 |
|  | AVGMOL | 0.247 | 75.30 | 12.28 | 12.42 |

Supplementary Data 1: Cell morphology and protein expression for *n* = 5,121,537 from 35,755 condensation grid boxes (Supplementary Data 2). <u>Uploaded to Dryad.</u>

Supplementary Data 2: Summary of cell morphology and protein expression for *n* =35,755 condensation grid boxes and associated data. <u>Uploaded to Dryad.</u>

